# Electrophysiological population dynamics reveal context dependencies during decision making in human frontal cortex

**DOI:** 10.1101/2022.10.11.511706

**Authors:** Wan-Yu Shih, Hsiang-Yu Yu, Cheng-Chia Lee, Chien-Chen Chou, Chien Chen, Paul W. Glimcher, Shih-Wei Wu

## Abstract

During economic choice, evidence from monkeys and humans suggest that activity in the orbitofrontal cortex (OFC) encodes the subjective values of options under consideration. Monkey data further suggests that value representations in the OFC are context dependent, representing subjective value in a way influenced by the decision makers’ recent experience. Using stereo electroencephalography (sEEG) in human subjects, we investigated the neural representations of both past and present subjective values in the OFC, insula, cingulate and parietal cortices, amygdala, hippocampus and striatum. Patients with epilepsy (n=20) reported their willingness to pay—a measure of subjective value—for snack food items in a Becker-DeGroot-Marschack (BDM) auction task. We found that the high frequency power (gamma and high-gamma bands) in the OFC positively correlated with the current subjective value but negatively correlated with the subjective value of the good offered on the last trial – a kind of temporal context dependency not yet observed in humans. These representations were observed at both the group level (across electrode contacts and subjects) and at the level of individual contacts. Noticeably, the majority of significant contacts represented either the present or past subjective value, but not both. A dynamic dimensionality-reduction analysis of OFC population trajectories suggested that the past trial begin to influence activity early in the current trial after the current offer was revealed, and that these two properties—current and past subjective values—dominate the electrophysiological signals. Together, these findings indicate that information about the value of the past and present rewards are simultaneously represented in the human OFC, and offer insights into the algorithmic structure of context-dependent computation during human economic choice.

## Introduction

Over the course of the last several decades, studies in macaque monkeys have come to define the electrophysiological representation of rewards and reinforcers (Schultz et al., 1997; Platt & Glimcher, 1999; Roitman & Shadlen, 2002; Fiorillo et al., 2003; Sugrue et al., 2004; Padoa-Schioppa & Assad, 2006, 2008; Kennerley et al., 2006; Rudebeck et al., 2017; Pastor-Bernier et al., 2021; Yang et al., 2022; for reviews, see Gold & Shadlen, 2007; Wallis, 2007; Kable & Glimcher, 2009; Kennerley et al., 2011; Padoa-Schioppa, 2011; Rudebeck & Murray, 2014; Klein-Flügge et al., 2022). This research has revealed that the firing rates of neurons in many brain areas encode a subjective estimate, the *subjective value*, of reward magnitude and type. Based on these extensive recordings, the broad topography of the network that represents reward-related value has now been well established in the macaque brain. Similar data are emerging for the rodent brain (Kepecs et al., 2008; Avila & Lin, 2014; Hanks et al., 2015; Constantinople et al., 2019; Gardner et al., 2019; Dabney et al., 2020; Lak et al., 2020), further extending our understanding of these important electrophysiological signals.

One key feature of this work in animals is that it has revealed the importance of context in the *subjectivization* of these reward-related signals. Monkey parietal cortex, for example, has been shown to encode a kind of spatial context-dependency (Louie et al., 2011; Rorie et al., 2010; Churchland et al., 2008). Monkey orbitofrontal cortex and dorsal anterior cingulate cortex, in contrast, appear to show a kind of temporal context-dependency, in which the recent history of rewards influences the electrophysiological representation of currently available rewards (Tremblay & Schultz, 1999; Padoa-Schioppa, 2009; Seo & Lee, 2009; Kennerley et al., 2011; Murray et al., 2014; Cavanagh et al., 2016). Closely related work has extended these findings to the rodents (Hocker et al., 2021). The importance of these findings, however, extends beyond the study of non-human animals because growing evidence suggests that these subjectivized representations seem to account for important idiosyncrasies and irrationalities observed in human choice behavior (Louie et al., 2013; Caplin & Dean, 2015; Khaw et al., 2017; Polania et al., 2018; Woodford, 2020; Webb et al., 2021).

At a neurobiological level, functional magnetic resonance imaging (fMRI) studies in humans have also linked animal-based studies of the subjective value network to our understanding of the human brain. While the blood oxygen level dependent (BOLD) signal measured by fMRI is quite distinct from electrophysiological measurements, many fMRI studies now show clear evidence of a subjective value network (Levy & Glimcher, 2012; Bartra et al., 2013; Clithero & Rangel, 2014) much like the one observed electrophysiologically in animals. These human fMRI studies indicate that subjective value representations in humans are much like those recorded from animal brains can be observed in the BOLD signal. Interestingly however, BOLD signal maps of subjective value in humans do not always agree with the electrophysiological maps developed in animals. In the parietal and orbitofrontal cortices, for example, very few studies using fMRI have identified the subjective value signals so often seen in non-human primate brains (though see Kahnt et al., 2014 for an example exception).

When considering fMRI data, however, much less information is available about the role of context in the human neural representation of reward. Although a tremendous amount of behavioral evidence identifies context-dependency as a critical factor that shapes both human and animal choice behavior (e.g., Kahneman & Tversky, 1979; Belke, 1992; Pompilio & Kacelnik, 2010; Zimmermann et al., 2018; Lin et al., 2020), few studies exist that localize context dependency in the human brain. Unlike in macaques, the evidence is unclear in humans whether and how the orbitofrontal cortex participates in context-dependent valuation (e.g., Elliot et al., 2008; Cox & Kable, 2014; Palminteri et al., 2015). This lack of clarity may reflect either a limitation of the technology, or a species difference. While there are some examples of context dependent responses in humans, it seems likely that the spatial and temporal scale at which fMRI operates and the nature of the BOLD signal itself, has made it extremely difficult to extract clear evidence of either spatial or temporal context dependency using that technology.

In this report we sought to achieve three goals aimed at addressing these gaps between our understanding of human and animal representations of reward and reinforcement. First, and most importantly, we sought to determine whether human intracranial electrophysiological signals encoding rewards show a clear and ubiquitous context dependency, as has been observed in animals. To that end, we focused the inquiry on temporal context dependency and sought to gather evidence indicating whether or not the recent history of rewards influences the electrophysiological representation in reward-encoding areas of the human brain. Second, we sought to perform this search at the single electrode (contact) and within-subject levels, which might allow us to overcome some of the limitations faced by region-of-interest based human intracranial electrophysiology studies. While averaging across subjects and electrode contacts has proven valuable in previous studies, animal research suggests that while the averaged signal may encode a property like reward value, this representation may be non-uniformly distributed from micro-site to micro-site. We hypothesized that by analyzing data at the single contact level, many of the important features which have never before been examined in humans could be assessed. Third, by analyzing data at the single contact level and transforming all recording sites to a standard anatomical reference, we hypothesized that it might also be possible to assess the spatial distribution of reward network signals at a very fine-grained level of analysis, as is common in animal research but has not yet been regularly undertaken in human studies.

The study reported here was thus designed to allow us to examine several subareas of the frontal and orbital cortex at relatively high resolution, as well as providing data at areal levels of analysis throughout several nodes of the reward value-network; the amygdala, hippocampus, striatum, insula, cingulate cortex, and parietal cortex. Our work extends critical earlier work by Pessiglione and colleagues (Lopez-Persem et al., 2019) and Hsu and colleagues (Saez et al., 2018) who were among the first to bridge the gap between human and animal research. Recording field potentials from human patients performing both rating and choice tasks, these authors provided the first electrophysiological confirmation that subjective value representations arise in the human brain in a manner very similar to what has been observed in the monkey.

Here we report the use of stereo electroencephalography (sEEG) to record neural activity in human epileptic patients (n=20) performing an incentive compatible valuation task known to induce temporal context dependency at the behavioral level in humans (Khaw et al., 2017). Our data show some of the first evidence for neurobiological temporal context-dependent value computations in humans. We observe this context dependency in a number of subregions of the orbitofrontal cortex. High-gamma activity (80-150 Hz)—thought to aggregate heterogeneous neuronal activity near the recording site (Buzsáki et al., 2012; Rich & Wallis, 2017)— represents both the subjective value of the present reward under consideration and the subjective value of the reward in the previous trial. The same patterns of correlation also arise in the gamma band (30-80 Hz). Our single-contact analysis reveals that at the level of gamma and high-gamma band signals, statistically significant single contacts encode *either* subjective value or temporal context, with only a few contacts encoding both: The orbitofrontal cortex carries context-dependent subjective value signals that seems to be built up from single contacts that encode either a subjective value signal that is only weakly influenced by temporal context or a clear and significant context-setting signal that is dissociable from the current reward. We found that context-dependent signals were more robustly observed in the central and medial orbitofrontal cortex than the lateral orbitofrontal cortex. In other brain areas we examined, the hippocampus and insula also carried these signals at the level of activity averaged across contacts *and* at the single contact level. Our single contact mapping data revealed that, as in monkey data (Rich & Wallis, 2016), high-frequency activity in only about 30% of recording sites carry statistically significant subjective value signals, and these sites are found to be distributed throughout each of the fronto-cortical and subcortical areas we examined. As in monkeys, not all locations within an area encode subjective value, and the locations which do are not apparently clustered but rather appear distributed throughout a given subarea. Finally, we adapted advanced dimension-reduction methods (Mante et al., 2013) developed to analyze neuronal population activity in macaques to characterize the population responses in humans. As in the monkey we found that activity in the human orbitofrontal cortex describes a spatial trajectory as a decision is being made, and that this spatial trajectory describes the time-course of human decision-making, including context dependency, just as it does the time course of monkey decision-making.

These results paint a novel and detailed picture of subjective value-related electrophysiological signals in the human brain at both the population and single-contact level. While broadly confirming previous findings from fMRI, they add to our understanding by providing some of the first evidence for context-dependent subjective value representations in the human brain, and the first multi-unit trajectory for a human decision in a frontal informational space. Perhaps unexpectedly, these data suggest that context-setting signals may be more patchily distributed than has been previously suspected, at least at the scale of sEEG. These findings are, it should also be noted, broadly compatible with at least some computational models of how context dependency arises in the subjective-value network (e.g., Glimcher, 2022).

## Results

In order to obtain behavioral measures of subjective value at the single-trial level, the subjects performed a version of the Becker-DeGroot-Marschak (BDM) auction task— a standard incentive-compatible paradigm used to elicit subjective value (Becker et al., 1964). On each trial, the subjects saw an image of a snack food item presented on a computer screen and had to indicate the maximum amount they were willing to pay for the snack food item (Fig. 1A). Here the amount is a measure of the subjective value for the food reward.

**Figure 1.**
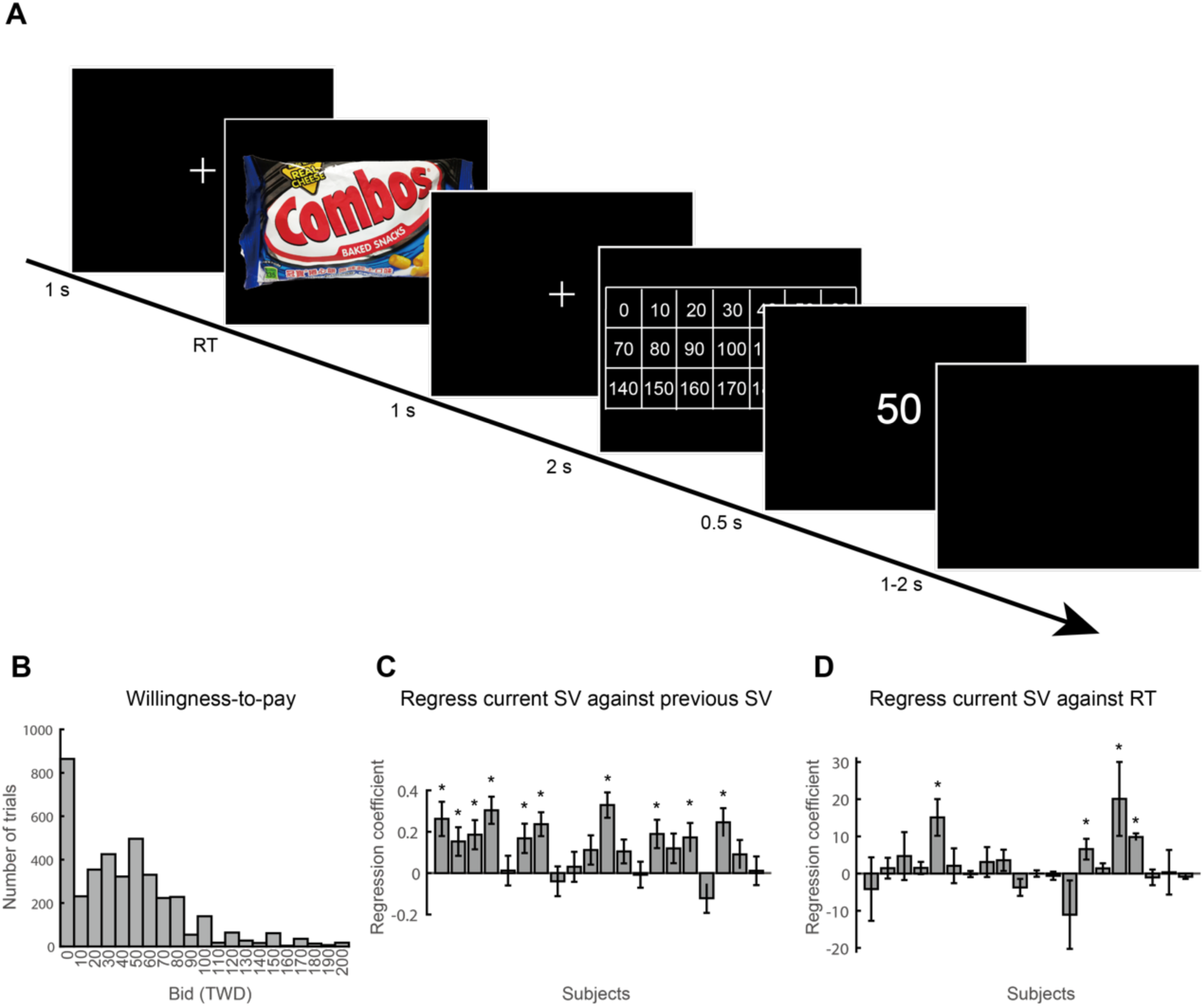
Experimental design and behavioral results. **A.** Trial sequence of the Becker-DeGroot-Marschak (BDM) auction task. On each trial, the subjects faced a snack food item and had to indicate their willingness to pay for that item. Subjects first pressed the left button on the mouse to signal that they were ready to indicate their willingness-to-pay. A matrix that indicated possible prices, from 0 to 200 cents, in 10 cent increments, would then appear on the computer screen. The subjects’ task was to use the mouse cursor to point to and click on the number closest to their maximum willingness-to-pay. **B.** The distribution of willingness-to-pay (in New Taiwan Dollars, TWD) from all subjects on all food items. **C.** The impact of the bid offered by the subject on the previous trial on willingness-to-pay on the current trial. Note that sequential items were selected randomly and in an uncorrelated manner for presentation. For each subject, we regressed their willingness-to-pay—a measure of subjective value (SV)—in the current trial against the willingness-to-pay in the previous trial. Here we plot the regression coefficient of the willingness-to-pay in the previous trial. In the majority of subjects, willingness-to-pay in the current trial was positively correlated with the willingness-to-pay in the previous trial. **D.** The relationship between the willingness-to-pay and response time (RT). For each subject, we regressed their willingness-to-pay against the response time in the trial. We plot the regression coefficient of the response time. In the majority of subjects, there was no relation between willingness-to-pay and response time. * indicates *p*<0.05.

### Bidding behavior

We found several interesting features in the subjects’ willingness-to-pay. First, across all subjects, the distribution of willingness-to-pay appeared to be positively skewed (Fig. 1B). About 23% of the trials across all subjects were zero bids. Second, the willingness-to-pay in a trial was significantly affected by the willingness-to-pay in the previous trial. The larger the subjects bid in a trial, the larger that she or he tended to bid in the next trial (Fig. 1C) even though sequentially presented rewards were uncorrelated in preceding BDM-bid values, indicating a temporal context dependency in bids. Third, we found no relationship between response time (how long it took the subjects to place the bid) and the amount of their willingness-to-pay (Fig. 1D). The distribution of individual subjects’ data on the willingness-to-pay and reaction time can be found in the Supplement (Supplementary Figs. 1 and 2). Individual subjects’ plots on the willingness-to-pay of the current trial against that of the previous trial and on the willingness-to-pay against response time can also be found in the Supplement (Supplementary Figs. 3 and 4).

### High-gamma activity in the orbitofrontal cortex represents past and present subjective value

In the orbitofrontal cortex (OFC), we collected stereo electroencephalography (sEEG) signals from a total of 166 electrode contacts in 20 subjects (Fig. 2A). The sEEG preprocessing pipeline can be found in the Supplement (Supplementary Fig. 5). Across these OFC contacts, we found that high-gamma (80-150 Hz) activity represented both the current subjective value (the willingness-to-pay of the current trial) and the past subjective value (the willingness-to-pay of the snack food item in the previous trial). Subjective-value representations were seen at both the group level (Fig. 2B) and at the individual-contact level (Fig. 2CD). At the group level, the mean time course of regression coefficients (averaged across all OFC contacts) showed that high-gamma activity positively correlated with the current subjective value, but negatively correlated with the previous subjective value (*p*<0.05, familywise error corrected for multiple testing across time points; see Permutation test in Methods for details).

**Figure 2.**
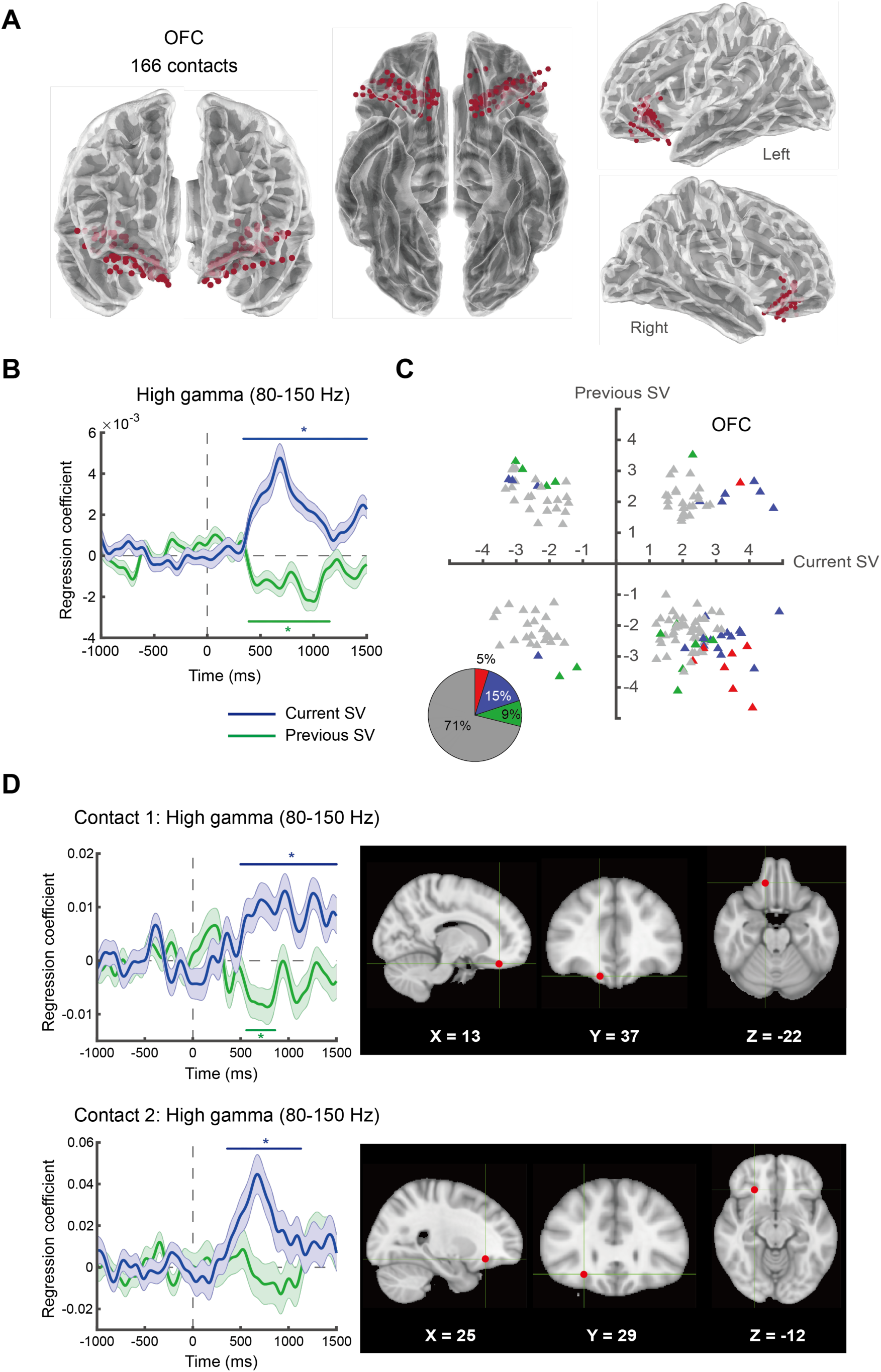
High-gamma activity (80-150 Hz) in the human OFC represents the subjective value of food rewards. **A.** Location of OFC electrode contacts. We collected data from a total of 166 electrode contacts in the OFC from 20 subjects. **B.** Subjective-value representations across OFC contacts. Here we plot the mean time course (averaged across all OFC contacts) of regression coefficients for the subjective value of the current trial (current SV, in blue) and the subjective value of the previous trial (previous SV, in green). **C**. Subjective-value representations from individual OFC contacts. We plot the *t* statistic of the current subjective value against that of the previous subjective value separately for each contact. Each data point represents a single contact. For each contact, we select the most significant time point according to the threshold-free-cluster-enhancement (TFCE) statistic, and plot the corresponding *t* statistic, separately for the current subjective value and previous subjective value. Since the *t* statistics come from the most significant time points, the data points in the graph are biased away from zero. Individual contacts that significantly represent the current subjective value, previous subjective value, or both are shown in blue, green, and red respectively. Individual contacts that neither represented the current nor the previous subjective value are shown in gray. The pie chart shows the proportions of contacts belonging to each of the categories described above. **D**. Results from two representative OFC contacts. Error bars represent ±1 standard error of the mean. Coordinates are in Montreal-Neurological-Institute (MNI) space. * indicates *p*<0.05 (familywise error corrected) using a permutation test based on the threshold-free cluster enhancement (TFCE) statistic. Colored (blue or green) horizontal lines indicate the time points with p<0.05 (familywise error corrected).

At the individual-contact level, we found that about 30% of the OFC contacts showed significant subjective-value representations (*p*<0.05, familywise error corrected at each contact) (Fig. 2C). The scatter plot (Fig. 2C) of *t* statistics according to the maximum threshold-free-cluster-enhancement (TFCE) statistic (Smith & Nichols, 2009; Winkler et al., 2014) revealed that the majority of significant OFC contacts cluster in the fourth quadrant, suggesting a positive correlation with the current subject value and negative correlation with the previous subjective value. This result is consistent with the group-level results (Fig. 2B). Among the significant OFC contacts, about half of them significantly represented the current subjective value, 29% of the contacts represented the previous subjective value, and 16% represented both the current and previous subjective value. In other words, a majority of the significant contacts represented *either* the current or the previous subjective value, but not both. Data from two representative contacts are also shown (Fig. 2D): One contact shows significant positive correlation with the current subjective value, and negative correlation with the previous subjective value. The other contact exhibits only a positive correlation with the current subjective value.

The presence of the previous subjective-value representation is consistent with the view that the OFC is sensitive to the temporal context of experience (Tremblay & Schultz, 1999). It is also consistent with the results in monkey OFC (Padoa-Schioppa, 2009; Kennerley et al., 2011) and with the view that the OFC implements a divisive-normalization algorithm to compute relative subjective value which we measure here using a simple linear regression (Yamada et al., 2018).

### Robustness of subjective-value representations in the OFC

To examine the robustness of the subjective-value representations, we performed four additional analyses to rule out potential confounds. First, we examined whether the results could have been driven by collinearity between the current and previous subjective value, since in most subjects the stated current subjective value positively correlated with the stated subjective value in the previous trial (Fig. 1C). To examine this possibility we carried out a regression analysis in two steps (GLM-2 in Fig. 3B). In the first step, we regressed high-gamma activity against the current subjective value. In the second step, we used the residuals from the first step as data and regressed them against the past subjective value. The results still indicated that the high-gamma activity in the OFC positively correlated with the current subjective value and negatively correlated with the past subjective value (Fig. 3B)—consistent with the original model (GLM-1 in Fig. 3A). We also performed the two-step analysis in the reversed direction and the results were identical (See Supplementary Fig. 6).

**Figure 3.**
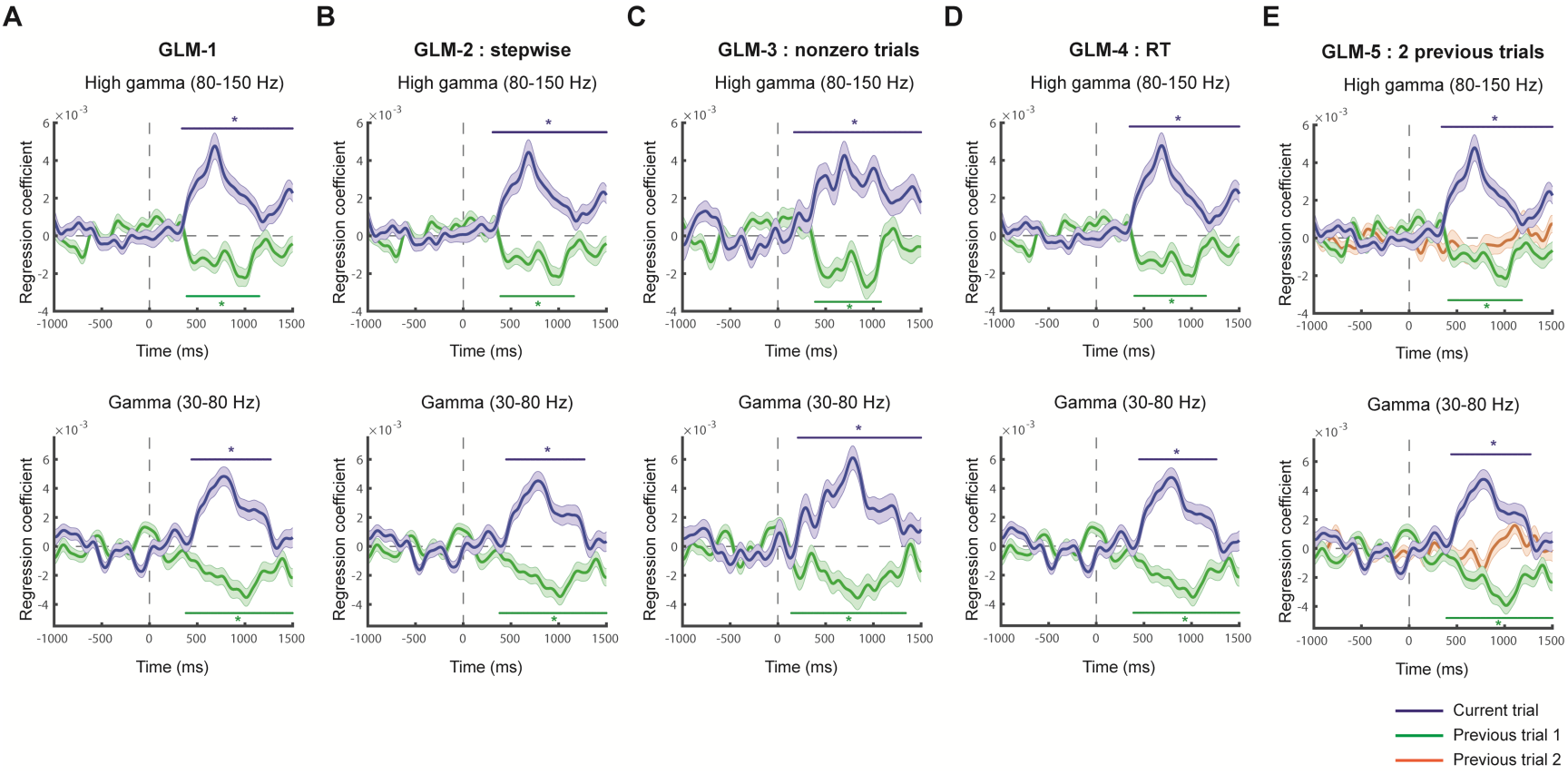
Testing the robustness of subjective-value representations in the OFC. We performed four different General-Linear-Modeling (GLM) analyses to examine the robustness of subjective-value representations in the OFC depicted in column A. The GLMs were implemented for both the high-gamma (80-150 Hz; top row) and gamma activity (30-80 Hz; bottom row). **A. GLM-1.** This is the original model where brain activity was regressed against the current and the previous subjective value. The high-gamma graph was identical to that shown in Fig. 2B. **B. GLM-2.** This analysis was performed in two steps. In the first step, we regressed brain activity against the current subjective value. In the second step, we used the residuals from the first step and regressed them against the previous subjective value. **C. GLM-3.** In this model, we only included trials where the subjects’ willingness-to-pay were not zero in the analysis. The model was identical to GLM-1. **D. GLM-4.** This regression model is identical to GLM-1 except that the subjects’ response time (RT) in the current trial was added as a nuisance regressor to the model. **E. GLM-5.** In this model, we added the subjective value of the option encountered two-trials back in addition to the subjective value of the current trial and the previous trial. * indicates p<0.05 (familywise error corrected) using permutation test based on the threshold-free cluster enhancement (TFCE) statistic. Colored (blue or green) horizontal lines indicate the time points with *p*<0.05 (familywise error corrected).

Second, we examined whether subjective-value representations can be affected by the zero-bid trials, as these trials represented about 23% of the total trials gathered across subjects (Fig. 1B). In GLM-3 we therefore excluded the zero-bid trials and only included the non-zero bid trials in the analysis. The results were again consistent with the original model (Fig. 3C). Third, we examined whether response time might somehow interact with subjective-value in a way that altered the results found in GLM-1. Therefore, in GLM-4, in addition to the current and previous subjective value as regressors, we added the subjects’ current-trial response time as a regressor to the model. Again, the results (Fig. 3D) were consistent with the original model (GLM-1 shown in Fig. 3A).

Fourth, we extended the original model by adding the subjective value obtained two trials previously as a regressor so as to examine whether the results would be consistent with the original results and to also examine the impact of the two-trials back bid on the OFC activity. The results were consistent with the original findings (Fig. 3E). However, we note that we found no significant impact of the subjective value in the two-trial back on the OFC activity, for both the high-gamma activity (upper graph in Fig. 3E) and gamma activity (bottom graph in Fig. 3E). While this almost certainly reflects a power issue, we are unable to conclude from this result whether or not the impact of previous trials extends back beyond one previous trial. Finally, we noticed that the patterns of subjective-value representations were remarkably similar between the high-gamma (80-150 Hz) activity (top row in Fig. 3) and the gamma (30-80 Hz) activity (bottom row in Fig. 3) across these five different regressions. This suggested that activity in a broad frequency range (from 30 to 150 Hz) represented the subjective value of food rewards in a similar fashion. Together, through these robustness checks support the conclusions that OFC electrophysiological activity in humans encodes both the subjective-value of the currently offered option and the subjective-value of at least one previously considered option.

### Subjective-value representations in different OFC subregions

We next examined whether patterns of subjective-value representations differed between the three major subregions on the human OFC—the medial (area 14), central (areas 11 and 13), and lateral (area 47/12) OFC. In all three regions, the high-gamma activity significantly correlated with the current subjective value (Fig. 4 in blue). By contrast, not all regions showed significant representation of the subjective value encountered in the previous trial: both the medial and central OFC significantly and negatively correlated with the previous subjective value, but not the lateral OFC (Fig. 4 in green). At the individual-contact level, the majority of contacts that significantly represented the current subjective value (blue triangles) showed positive correlation with the current subjective value. The results on the previous subjective value, at the individual-contact level, were less consistent across the three regions. In the central OFC, significant previous-value contacts tended to show a negative correlation with previous subjective value. In the medial and lateral OFC, this tendency was less obvious.

**Figure 4.**
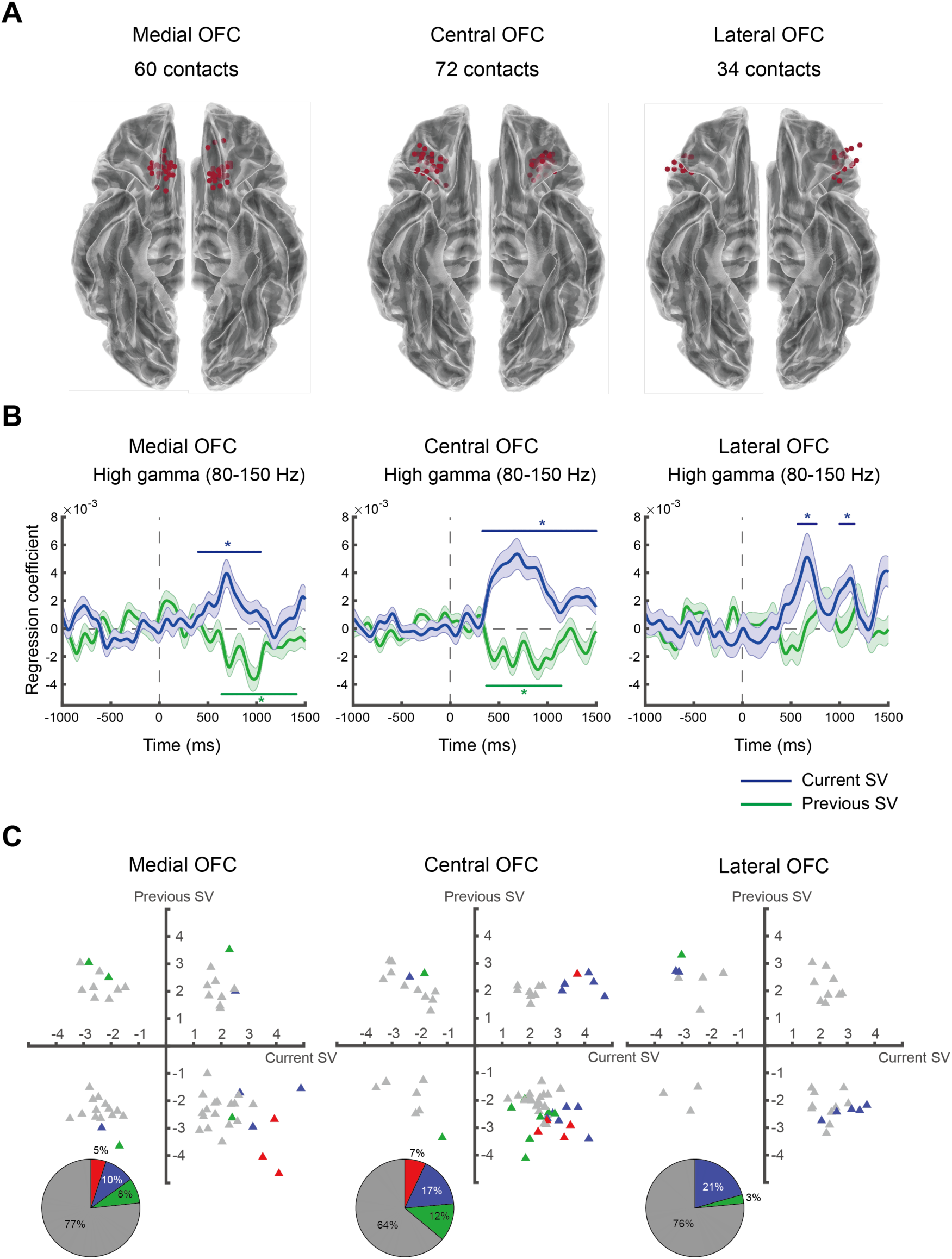
Subjective-value representations in the three major regions of the OFC. **A.** Electrode contacts in the medial (left), central (middle), and lateral OFC (right). **B.** High-gamma activity. Average time course of regression coefficients for the current subjective value (in blue) and previous subjective value (in green) in the medial, central, and lateral OFC. * indicates p<0.05 (familywise error corrected) using permutation test based on the threshold-free cluster enhancement (TFCE) statistic. Colored (blue or green) horizontal lines indicate the time points with *p*<0.05 (familywise error corrected). **C.** Subjective-value representations in individual OFC contacts. Conventions are the same as described in Fig. 2C.

### Cross-frequency representations of subjective value

Surprisingly, in the OFC, we not only found significant subjective-value representations in the high-gamma and gamma activity bands (Fig. 3), but also in the activity of lower frequencies (Fig. 5). The two-dimensional heatmap (Fig. 5A) plots the *z* statistic for the regression coefficients (across all electrode contacts across all subjects) in the time-frequency space for the current subjective value (left graph in Fig. 5A) and for the previous subjective value (right graph in Fig. 5A). The colors in the maps reveal the encoding directions of subjective value—orange for positive correlation with the subjective value, blue for the negative correlation, and green for non-significant results. It is evident that, after stimulus onset (indicated by 0 on the horizontal axis), activity in the gamma and high-gamma band positively correlated with the current subjective value (the orange clusters in the left graph, Fig. 5A) but negatively correlated with the previous subjective value (the blue clusters in the right graph, Fig. 5A). Interestingly, the encoding patterns of subjective value were reversed in the low frequency bands. Activity in the beta (13-30 Hz), alpha (8-12 Hz), and theta (4-7 Hz) bands negatively correlated with the current subjective value (blue clusters in the left graph, Fig. 5A), but positively correlated with the previous subjective value (orange clusters in the right graph, Fig. 5A). These results are further summarized in the group-level time series plots of the regression coefficients (Fig. 5B). At the individual-contact level, scatter plots of the *t* statistic according to the maximum TFCE statistic are plotted in Fig. 5C. Noticeably, the low-frequency activity significantly represented a bias on the current subjective value before stimulus onset—before information about the current food item was revealed. We found that in the alpha band, these results were associated with two behavioral patterns: the variability of the bids and the correlation between the current subjective value and the previous subjective value. We found that, in part, such pre-stimulus representations were driven by the subjects who showed less variability in their bids and whose bids were more affected by the bid in the previous trial (Supplementary Figs. 7 and 8). In other words, the more the subjects relied on the previous bid, the greater the likelihood of significant pre-stimulus representations (a form of bias).

**Figure 5.**
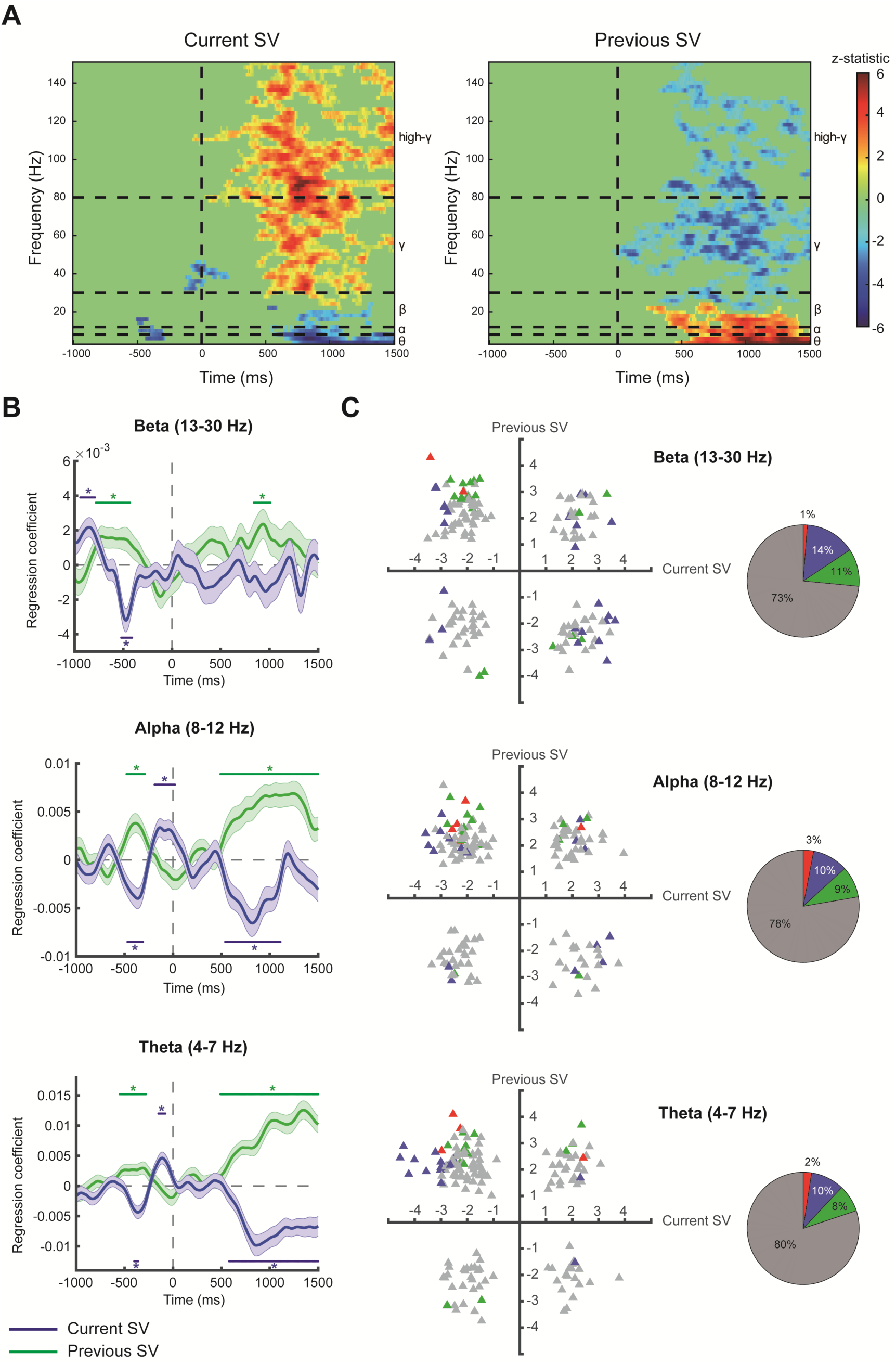
Cross-frequency representations of subjective value. **A.** Time-frequency representations for subjective value. The heatmap plots the *z* statistic of the regression coefficient for the current subjective value (left graph) and previous subjective value (right graph). Significant clusters in these two-dimensional maps were identified by contiguous points in the time-frequency space that survived multiple testing after with a familywise error rate of 0.05 according to the threshold-free-cluster-enhancement (TFCE) statistic. Significant positive correlation is represented by orange, significant negative correlation is indicated by blue. Non-significant results are coded by green. **B-C.** Results from activity in the beta, theta, and alpha bands. **B.** Average time course of regression coefficients for the current subjective value (blue) and previous subjective (green) in the beta (13-30 Hz, top graph), alpha (8-12 Hz, middle graph), and theta (4-7 Hz, bottom graph) activity. **C.** Subjective-value representations on individual OFC contacts in the beta (top graph), alpha (middle graph), and theta activity (bottom graph). Conventions are the same as described in Fig. 2C. * indicates p<0.05 (familywise error corrected) using permutation test based on the threshold-free cluster enhancement (TFCE) statistic. Colored (blue or green) horizontal lines indicate the time points with *p*<0.05 (familywise error corrected).

### Subjective-value representations in other brain regions

We also examined subjective-value representations in several other subcortical (Fig. 6) and cortical regions (Fig. 7). The subcortical regions included the amygdala, hippocampus, and striatum. At the group-level, all of these regions except the striatum showed significant subjective-value representations (middle graph in Fig. 6). Both the amygdala and the hippocampus significantly represented the current subjective value, while the hippocampus also represented the previous subjective value. In the hippocampus, at the individual-contact level, the majority of the significant contacts seemed to cluster in the fourth quadrant, suggesting a positive correlation with the current subjective value and a negative correlation with the previous subjective value—a result consistent with our findings in the OFC. In the amygdala, 90% of the contacts were non-significant, even though the group-level results indicate significant current-value representation in the area. To summarize, even though all three subcortical regions showed evidence for subjective-value representations, only the hippocampus showed significant representations for both the current and previous subjective value, at the group level and at the level of individual contacts.

**Figure 6.**
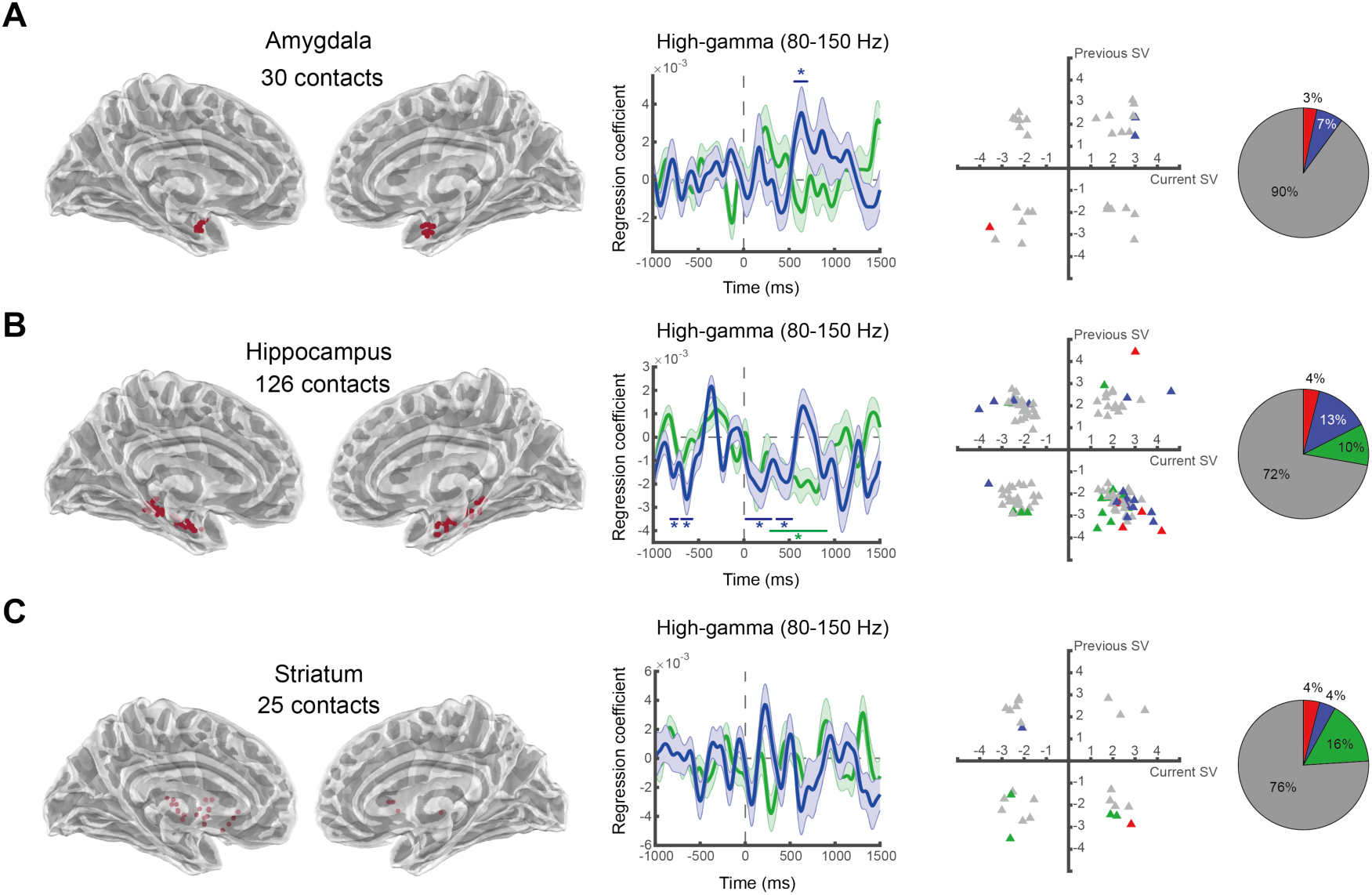
Subcortical representations for subjective value. High-gamma activity in the amygdala, hippocampus, and striatum. Conventions are the same as described in Figure 2. **A.** Amygdala. **B.** Hippocampus. **C.** Striatum.

**Figure 7.**
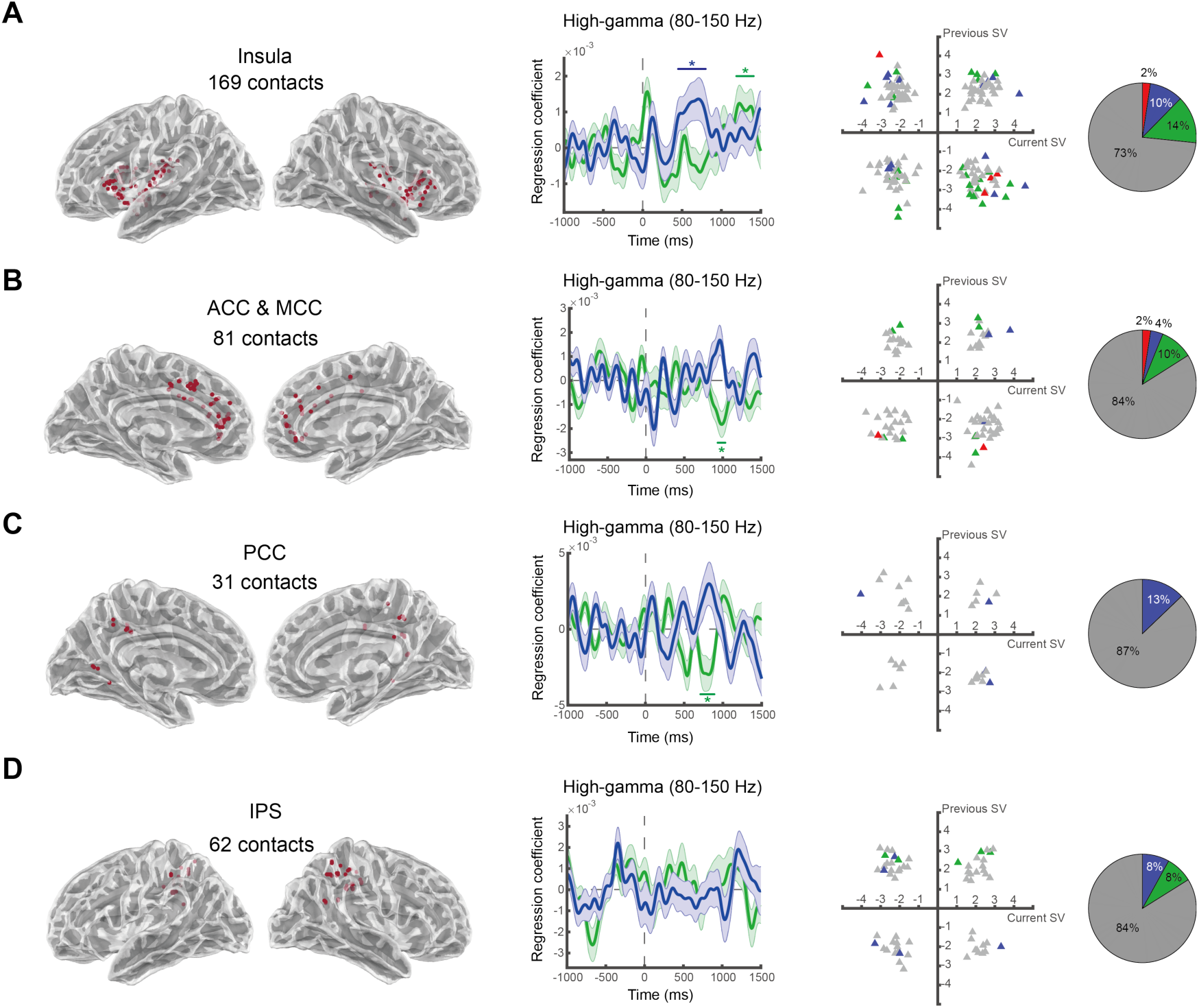
Cortical representations for subjective value. High-gamma activity in the insula, cingulate cortex, posterior cingulate cortex, and intraparietal sulcus. Conventions are the same as described in Figure 2. **A.** Insula. **B.** Anterior cingulate and midcingulate cortex (ACC and MCC). **C.** Posterior cingulate cortex (PCC). **D.** Intraparietal sulcus (IPS).

In our analysis of recordings from cortical regions other than the OFC—including the insula, the anterior cingulate and midcingulate cortex (ACC and MCC), posterior cingulate cortex (PCC), and the intraparietal sulcus (IPS)—we found that, at the group level, all but the IPS significantly represented the subjective value. The insula significantly represented both the current and the previous subjective value, while the ACC-MCC, and PCC represented the previous subjective value. At the individual-contact level, about 85% of contacts in the ACC-MCC, PCC, and IPS were non-significant, while about 30% of the contacts in the insula were significant in either representing the current or the previous subjective value. The proportion of significant contacts in the insula (27%) was identical to the OFC (29%, Fig. 2C) and hippocampus (28%, Fig. 6B). In summary, while there was evidence for subjective-value representations in these four cortical regions, only the insula showed prominent representations for both the current and previous subjective value, as these representations were observed at the group level and at the level of individual contacts in our valuation task.

### OFC activity trajectories in value space

To further understand how the OFC as a whole dynamically represents subjective value and context, we performed two final complementary analyses, one based on a principal component analysis (PCA) and the other based on a regression subspace analysis (Mante et al., 2013; Aoi et al., 2020).

For each electrode contact, we first computed the mean high-gamma activity timeseries (across all trials) and subtracted it from each individual trials’ timeseries. Second, we sorted trials according to the subjective value of the current trial (high or low, median spilt) and the subjective value of the previous trial (high or low, median split). The medians we used were subject-specific. This resulted in a 2 (current subjective value magnitude) × 2 (previous subjective value magnitude) design and a total of four conditions. Third, we gathered the activity timeseries across all OFC contacts according to condition (Fig. 8A). Each condition is represented by a two-dimensional matrix where each row represents the timeseries—from 1s before stimulus onset to 1.5s after stimulus onset—of a single OFC electrode contact. Fourth, we stacked up the four two-dimensional matrices (one for each condition) and performed a single principal component analysis (PCA) across all conditions. The PCA allowed us to identify the dimensions in the neural state space that captured the most variance in the OFC high-gamma activity. Here the neural state space is a *N_contacts_*-dimensional space where *N_contacts_* is the number of electrode contacts in the OFC (*N_contacts_* = 166). This analysis approach combines two advantages in our dataset—the idiosyncratic preferences of different individuals and a large number of electrode contacts across participants. In other words, by using the individual-specific medians to categorize trials into different conditions, we preserved the individual-specific preference information in this population-level, across-subjects analysis.

**Figure 8.**
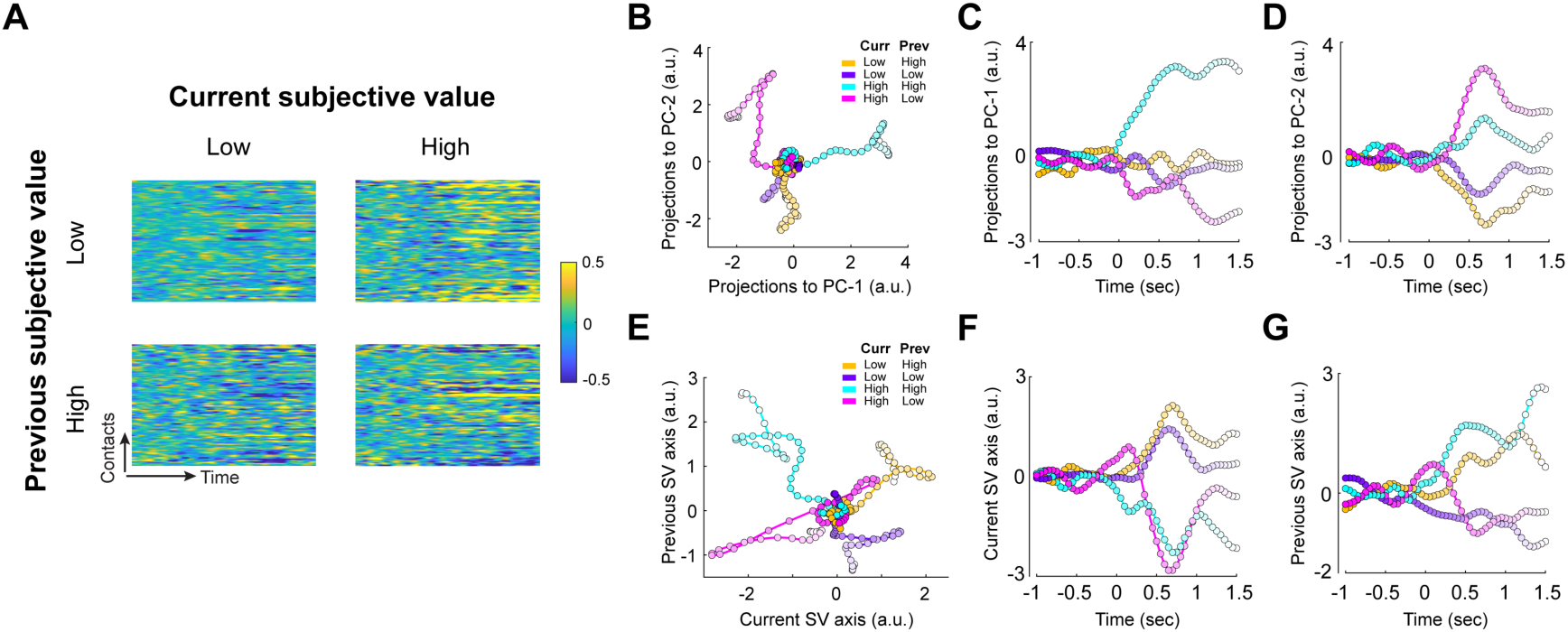
State-space analysis reveals OFC activity trajectories in value space. **A.** OFC activity sorted by condition. We sorted trials into four different conditions according to the magnitude of the current subjective value (high or low, median split for each subject separately) and the previous subjective value (high or low, median spilt for each subject separately). Each graph summarizes the average OFC activity timeseries of 166 electrode contacts. Each row represents the timeseries—from 1 s before stimulus onset to 1.5 s after stimulus onset—of a single OFC contact. **B-D.** Principal component analysis (PCA) of all trials massed together reveals distinct, condition-specific activity trajectories and subjective-value axes. **B.** Activity trajectories for subsets of trials all used in the PCA are plotted and color-coded with respect to the four conditions in the space of the first and second principal components. The horizontal and vertical axes represent the projections of OFC population activity onto the first principal component (PC-1) and the second principal component (PC-2). Time is indexed by the darkness of the colors, with the starting time point (1 s before stimulus onset) being the darkest and the end time point the lightest. Each data point in the trajectories represents a 50-ms time point. **C.** PC-1 captures the subjective value from the previous trial. Here we plot the activity trajectories on the PC-1 axis as a function of time. High previous subjective value: cyan and yellow; low previous subjective value: magenta and purple. **D.** PC-2 captures the subjective value of the current trial. Here we plot the activity trajectories on the PC-2 axis as a function of time. High current subjective value: magenta and cyan; low current subjective value: yellow and purple. **E.** Activity trajectories are plotted in the space constructed of the current subjective-value axis (horizontal axis) and the previous subjective-value axis (vertical axis). Color codes are identical to **8B**. **F.** The current subjective-value axis distinguishes high (magenta and cyan) from low (yellow and purple) current subjective value. Here we plot the activity trajectories on the current subjective-value axis as a function of time. **G.** The previous subjective-value axis distinguishes high (cyan and yellow) from low (magenta and purple) previous subjective value. Here we plot the activity trajectories on the previous subjective-value axis as a function of time.

We then projected the activity timeseries from each of the four groups of trials (current subjective value magnitude: high and low × previous subjective value magnitude: high and low) separately onto the first principal component (PC-1) and the second principal component (PC-2) and plotted a temporal trajectory of the population activity during each of these four groups of trials (Fig. 8B). We found that each of the four conditions had very unique trajectories, suggesting that the top two principal components—the factors that accounted for the most variance in the OFC high-gamma activity—strongly and orthogonally capture information about current and past trial subjective value. The four activity trajectories start from a common origin (corresponding to 1s before stimulus onset) and then diverge in four different cardinal directions. These directions revealed that the first principal component appears to capture the subjective value observed in the previous trial (Fig. 8C), separating high previous subjective value (yellow and cyan) from low previous subjective value (purple and magenta). The degree of separation appears to depend on the current subjective value, with stronger separation between high and low previous subjective value when the current subjective value was high (cyan and magenta). The onset of this stronger separation emerged early, right after the stimulus onset (0s mark). The second principal component appears to capture information in the population about the magnitude of the subjective value observed in the current trial (Fig. 8D), separating high current subjective value (cyan and magenta) from low current subjective value (yellow and purple). The emergence of this separation also appeared to be early, approximately 200 ms after stimulus onset. These patterns were consistently observed under different data-smoothing parameters (Supplementary Fig. 9). This unbiased PCA analysis thus seems to support the notion that, at least when human subjects are performing a valuation task, much of the variability in activity observed in the OFC encodes the current value and context, with context information seeming to have a greater overall impact on the data variance.

Interestingly, the regression subspace analysis adapted from Mante et al. (2013 revealed a similar pattern of trajectories (Fig. 8E-G) as those observed in the PCA (Fig. 8B-D). The regression subspace analysis aims to reveal OFC activity in a value-specific low-dimensional subspace that captures the across-trial variance due to the subjective value of the current trial, and the subjective value of the previous trial. The analysis consisted of two steps. First, we denoised the OFC population activity (using the first 12 PCs from PCA accounting for 78.59% of the neural variance across electrode contacts). Second, we projected the denoised population activity onto two orthogonal axes—the axis of the current subjective value (horizontal axis) and the previous subjective value (vertical axis)—that were defined based on the regression coefficients of subjective value in GLM-1 (See Methods for details). The trajectories from the four different conditions again started at the same origin and quickly established their unique paths after stimulus onset in a manner highly similar to the one observed in the first two components of the raw PCA. As expected then, the current subjective value regression axis could easily distinguish the current subjective value (Fig. 8F), separating high current subjective value (cyan and magenta) from the low current subjective value (purple and yellow). The emergence of this separation appears to begin within 500ms after stimulus onset. The previous subjective value regression axis could easily distinguish the previous subjective value (Fig. 8G), separating the high (yellow and cyan) from the low (purple and magenta) previous subjective values. Finally, we found that these patterns remained when we varied the number of PCs for activity denoising (from 2 to 20 accounting for 39.89% and 89.83%; See Supplementary Fig. 10) and when no data smoothing was applied (Supplementary Fig. 11).

In summary, these two analyses reveal the dynamics of OFC activity in representing the subjective value and temporal context, and highlight the fact that context is a major source of organized patterns of activity in the OFC. At least while subjects are performing valuation tasks, value and context seem to be the major determinants of activity patterns in this region.

## Discussion

In this study, we used stereo electroencephalography (sEEG) in human epileptic patients to investigate the electrophysiological representation of subjective value and context, in humans—fundamental building blocks in the theory of decision making widely studied in non-human animals. Our data show that, as observed previously in animals, human subjective value signals show strong evidence of context dependency. Indeed, our PCA results suggest that context is an even more significant determinant of activity pattern than simple value. Previous work has indicated that the human orbitofrontal cortex (OFC) represents the subjective value of rewards under immediate consideration: Gamma (30-80Hz) and high-gamma activity (80-150Hz) have both been shown to positively correlate with the subjective value of an offered reward (Lopez-Persem et al., 2019). We found that these signals were negatively influenced by the magnitude of rewards offered on previous trials, a form of temporal context dependency that has been observed behaviorally in humans and physiologically only in non-human primates. To our knowledge, this is the first evidence of context-dependency in human electrophysiological signals encoding subjective value.

An important—and often less explored in previous studies—dimension of our data analysis was an analysis of signals at the single contact/electrode level. Many previous human intracranial studies have been forced to average across electrode contacts in order to report findings about a region of interest. The high quality of both our initial signals and the nature of our analytic pipeline, allowed us to both examine single-contact level data and region of interest averages. The single-contact level data allowed us to examine the local spatial distribution of both subjective value and context signals, independently, at the level of cortical subareas. Our data revealed that signals from many individual contacts in the lateral OFC encode a subjective value signal that is only weakly influenced by temporal context. In contrast, subjective value signals influenced by context were more common in the central and medial subregions of the orbitofrontal cortex. In a similar vein, we found that the hippocampus and insula carried subjective value signals strongly influenced by temporal context at the single contact level.

Our contact-by-contact data also indicates that even in areas known to carry robust subjective value signals, only about 30% of the recording sites carry those signals (in a statistically significant way). The observed patchy distribution of subjective value signals in the human agrees well with parallel work in the monkey conducted with much finer electrodes gathering signals at the single neuron level.

### Value representations in the human and monkey OFC

There is now widespread agreement that subjective value representations are broadly distributed in the mammalian brain. A series of influential meta-analysis studies examining fMRI data from human subjects have focused attention on two key areas: the ventral striatum and the ventromedial prefrontal cortex. Activity in these two areas is now widely associated with decision-making and subjective value and measurements of these areas are now often used as direct tools for assessing subjective value in humans. Interestingly, however, the focus on these two *human* brain areas stands in contrast to extensive electrophysiological work on value representations in non-human primates and to a lesser extent in rodents. Multiple electrophysiological studies in animals have focused interest on the orbitofrontal cortex as critical source of subjective value signals in these species. This apparent mismatch between human and monkey data has been of some significance, and has raised the question of whether this mismatch reflects a technological difference between fMRI and electrophysiology or a species difference between humans and other mammals.

Recently, a small number of intracranial electrophysiological studies have begun to address this dichotomy by using electrophysiological tools to examine subjective value representations in humans. Studies on value-based decision making in humans have now been conducted using electrocorticography (ECoG) (Saez et al., 2018) and sEEG in human epileptic patients (Jenison et al., 2011; Mormann et al., 2019; Lopez-Persem et al., 2020). Since the high-frequency components (the gamma and high-gamma bands) of the sEEG and ECoG signals are believed to correlate tightly with the single unit activity (Siegel & König, 2003; Liu & Newsome, 2006; Berens et al., 2008; Belitski et al., 2008; Ray & Maunsell, 2011; Perge et al., 2014; Rich & Wallis, 2017), it should be possible to use these tools to search for the human analog of monkey OFC value signals. Lopez-Persem and colleagues (2020) for example, used a region of interest approach with sEEG recordings and showed unambiguous subjective value signals that were the human homologue of monkey value signals in that same region.

Our observations extend this earlier work from the level of a region of interest to the level of a single recording contact. This adds to a growing literature using sEEG or ECoG in humans that finds subjective-value representations in the medial (area 14) and central OFC (areas 11 and 13) in single-unit responses (Mormann et al., 2019), in gamma and high-gamma activity (Saez et al., 2018; Lopez-Persem et al., 2020), and in single-unit activity in the amygdala (Jenison et al., 2011). It should, however, be noted that the naming of the subdivisions of OFC are not consistent across studies. Here we have adopted the monkey-based convention used by Padoa-Schioppa and Cai (2011) and Wallis (2012) in describing the medial (area 14), central (areas 11 and 13), and lateral (area 47/12) OFC.

### Measuring subjective value: choice, willingness-to-pay, and liking ratings

Our current understanding of the subjective valuation of rewards in humans is based on three different methods for eliciting subjective value—choice tasks, the Becker-DeGroot-Marschak auction task where subjects indicate willingness-to-pay, and liking-rating tasks. To date, however, there has been little discussion of how these different elicitation methods might differentially impact the subjective-value signals measured physiologically.

While we employed the incentive compatible Becker-DeGroot-Marschak value elicitation method to assess a subject’s subjective valuation for food items (as in Jenison et al., 2011), most other studies have employed either the liking-ratings tasks under similar conditions (Mormann et al., 2019; Lopez-Persem et al., 2020) or asked the subjects to choose between different monetary lotteries (Saez et al., 2018). We chose the willingness-to-pay approach presented here for two reasons. First, unlike the liking-rating approach, with the BDM it is in the subjects’ best interest to provide their true valuation of the rewards and some research indicates that this yields more accurate estimates of subjective value at the behavioral level (Becker et al., 1964). Second, compared with choice tasks that offer 2 or more options, each trial presents only one object for evaluation, simplifying the interpretation of neural data to the representation of a single object.

Despite these advantages of the BDM approach, however, it must be acknowledged that the cognitive and motivational processes associated with each method are not identical. How might these different elicitation methods impact subjective valuation? It will be important for future investigations to enrich our understanding of the neural representations for subjective value by characterizing the similarities and differences between these different methods as they impact subjective value signals in the brain.

### Contextual Modulations of Subjective Value Representations

Numerous studies in non-human primates have made it clear that the subjective value signals in the OFC are strongly influenced by context. Padoa-Schioppa and colleagues (Padoa-Schioppa, 2009), for example, have shown that the firing rates of single monkey neurons are strongly adapted by temporal context. If animals face blocks of low value rewards, this has the effect of reducing the responses of OFC neurons. These non-human primate signals appear to be affected the by the range of subjective value experienced in the recent past (Simmons & Richmond, 2008; Padoa-Schioppa, 2009; Kobayashi et al., 2010; Kennerley et al., 2011). It remains an open question as to whether the human OFC activity exhibits this same property. Another important and open question is how these relative-value signals in OFC contribute to choice behavior. While studies had begun to show the effects of range on choice behavior (Zimmermann et al., 2018), the links between OFC relative-value signals and range-affected choice behavior remain unclear.

Our data show that recently experienced rewards influence the activity of human subjective value neurons in these same areas. We found that the reward delivered on the preceding trial effectively down-adapted the signal observed from the human OFC. This is a finding compatible with both the standard range adaptation models and with divisive normalization models, both of which are aimed at describing the influence of recent temporal context on OFC firing rates. We stress that the fact that our linear regression analysis suggests a subtractive relationship between current and previous reward should not be interpreted as specifically supporting a subtractive relationship in the neuronal data. Linear regression is constrained to always represent divisive relationships as subtractive. Were the true relationship divisive it would be expected to appear subtractive upon linear regression as used here. For this reason, we must be silent about the true form of the representation.

### Population-Level Analyses Using Dimension Reduction Approaches in Humans

Over the last decade there has been increasing interest in aggregating information from large populations of neurons in macaques and rodents into high dimensional datasets that can then be analyzed at the population level (Cunningham & Yu, 2014; Vyas et al., 2020; Urai et al., 2022). Of particular interest has been the extraction of the trajectories—in low-dimensional space—neuronal activity takes in these high dimensional data-spaces in valuation and decision-making tasks. In this report we extended those approaches to the study of human sEEG signals, for what we believe to be the first time. Our results both validate and extend this earlier work in non-human primates. First, our unbiased PCA revealed that during our task the population begins each trial at a common starting point and then evolves toward a representation whose primary properties are a representation of reward context and current offer, with the suggestion of context being an even larger signal than value. Our regression subspace dimensionality reduction analysis (Mante et al., 2013) further confirmed and extended this finding, revealing that OFC population dynamics formed distinct trajectories according to the subjective value and context setting signals. These trajectories serve as another point of contact with monkey data, reinforcing the similarities of these two species in the orbitofrontal cortex.

### Single Contact Measurements Reveal Site Heterogeneity

Previous monkey studies have also provided some sense of how subjective value and context signals are distributed in the monkey brain. While subjective value signals are observed robustly in areas like the monkey OFC, it is important to note that not all OFC neurons show these signals. Estimates from Rich and Wallis (2017) made in the monkey suggest that only about 30% of channels in the OFC carry a subjective value signal. They found that after stimulus onset, the peak percentage of OFC channels whose high-gamma activity represented expected reward size was about 20%. After a reward was delivered, the peak percentage was about 40% in representing the type of reward the animals received. Interestingly, we observed a similar result: in high-gamma activity, about 30% of our OFC electrode contacts showed evidence of either the subjective value signals, or the context setting signals (subjective value of the previous trial), or both. This seems to show excellent agreement between humans and monkeys.

Our examination of context dependency, however, does seem to suggest a difference between humans and monkeys. We observed that in some areas, like the central and medial OFC, individual contacts reflected either subjective value signals or context setting signals. Very few contacts represented both the subjective value and context setting signals. This is an observation that has not been widely reported in the monkey. While it will be important to confirm these findings, this does raise the possibility that human context setting signals may be distinctive in some way.

### Cross-frequency representations

Results from non-human primates also may shed light on how broad-band sEEG signals might be expected to behave. For example, it has been suggested that low-frequency activity in the alpha band may be involved in modulating inputs from task-relevant and task-irrelevant brain regions (Haegens et al., 2011). In studies of reward representations, An et al. (2019) found that in non-human primates, reward expectation increased single-unit firing rates in the primary motor cortex but decreased alpha (8-14 Hz) oscillatory power. Given that single-unit firing rates often positively correlate with high-gamma power (Ray & Maunsell, 2011), this finding suggests that the high-frequency power (e.g., high-gamma activity) in M1 may positively correlate with reward expectation, while low-frequency power, such as alpha power, negatively correlates with reward expectation. This is similar to what we found where the encoding directions for subjective value in the low-frequency activity were reversals of those in the high-frequency activity.

One potential explanation for the encoding directions of subjective value in the alpha band is the inhibition of the previous subjective value. Increase in alpha oscillations have been shown to reflect inhibitory activity in circuits associated with attention, perception, and working memory (Foxe et al., 1998; Jensen et al., 2002; VanRullen & Koch, 2003; Sauseng et al., 2005; Thut et al., 2006; Rihs et al., 2009; Kerlin et al., 2010; Jensen & Mazaheri, 2010). Hence, it is possible that the positive correlation of alpha activity with the previous subjective value reflects the inhibition of information about past subjective value when the subjects evaluate a snack food item in the current trial. Similarly, the negative correlation of alpha power with the current subjective value may reflect increased attention to the food item in the current trial.

Our findings also suggest the involvement of alpha oscillations in modulating visual attention during value-based decision making (Krajbich & Rangel, 2011). Previous fMRI studies found that value signals in the vmPFC are modulated by visual attention (Lim et al., 2013). An open question, therefore, is to investigate whether and how alpha oscillations in the OFC, and other value-related regions, change in a free-choice paradigm where different options are simultaneously presented and the subjects are free to look at these options before making a decision.

### Implications to the links between LFP and BOLD signals in value representations

Our results were consistent with the view that there is a close relationship between high-frequency oscillations (30 to 150 Hz) in local field potential (LFP) and the fMRI BOLD signals. Our results showed the involvement of gamma (30-80 Hz) and high-gamma (80-150 Hz) activity in the representation of subjective value. This frequency range (30-150 Hz) coincides with previous observations that “LFPs were often dominated by stimulus-induced and usually stimulus-locked fast oscillations in the range of 30-150 Hz” (Logothetis et al., 2001). Given that BOLD signals have been found to be better described by the LFP than by multi-unit activity (MUA), it is possible that the gamma and high-gamma findings here would be observed in the BOLD signals in the human OFC had there been the same spatial resolution or no signal loss due to the susceptibility artifacts in the BOLD signals in this brain region. In brain regions associated with subjective value representation that do not suffer BOLD signal loss, we would expect the gamma and high-gamma activity there to represent subjective value. Indeed, we found that many other brain regions also represented the subjective value in high-gamma activity. Among them, the insula and hippocampus stood out because evidence for the past and present subjective-value representations were found at the group level (averaged across electrode contacts) and at the level of individual contacts in those regions.

### Technical limitations of sEEG

An important limitation for any sEEG is the sparse and heterogeneous coverage of the electrodes. The decision about where to implant electrodes is, rightfully, an entirely clinical decision, but as a result the spatial coverage we can achieve is spare and heterogeneous, both within and across the subjects. This limitation poses challenges to both within- and between-subject (group-level) statistical inference and the interpretation of null results. On the one hand, null results can signal that a region is not involved in certain tasks or computations. On the other hand, the null results could be driven by sparse or inefficient coverage. In the context of our study, it is insufficient to conclude, for example, that OFC does not represent subjective value on a particular subject based on the null results from his or her OFC contacts. It is possible that his or her OFC contacts are not ‘in the right spot’—regions in the OFC that represent subjective value. One possible way to address this issue is the development of distributed, anatomically realistic source modeling of LFP data (e.g., Lin et al., 2021). Future studies need to explore this direction and to examine its feasibility and value in contributing to the interpretations of LFP signals in human sEEG experiments.

### Summary

Many of the behaviors in humans and animals are affected by the context of our recent experience. Characterizing the representations of context at the computational, algorithmic and neural implementation levels, therefore, is essential to understanding a wide array of cognitive functions. Using human intracranial electrophysiology, we found several distinct features of context-dependent representations for the subjective valuation of rewards. At the computational and algorithmic levels, temporal context— recent history of rewards—was represented in a manner predicted by existing models like divisive normalization and range adaptation. At the neural implementation level, we found that the current reward value and the context were represented by distinct electrode contacts in the orbitofrontal cortex, insula, and hippocampus. These findings suggest that contextual adaptation is implemented through distinct, large-scale neuronal populations that separately represent current and past information about reward value.

## Methods

### Participants

Twenty patients with drug-resistant epilepsy participated in this study (9 males; aged 16–51 years; average: 29.2 years). Patients had been implanted with multi-contact depth electrodes and were undergoing intracranial monitoring in order to identify seizure onset regions. Each patient was implanted with 7 to 14 electrodes. The decision to implant the electrodes and their location was driven solely by medical considerations. The study was approved by Taipei Veterans General Hospital Institutional Review Board. Informed consent was obtained from each patient before participation.

### Behavioral Task

The subjects performed a version of the Becker-DeGroot-Marschak (BDM) auction task during sEEG recording. They were asked to refrain from eating for at least two hours before the start of the experiment. Prior to the BDM task, the subjects received 200 New Taiwan Dollar (TWD) as an endowment to purchase food items.

The BDM task consisted of 8 blocks of 25 trials each. One hundred snack food items were introduced and each food item was presented twice in the experiment. In each trial, a snack food item was presented and the subjects were instructed to bid— his or her maximum willingness-to-pay—for the snack food item. A trial started with a fixation cross presenting in the center of the screen for 1 s. Following the fixation, a snack food item was presented on the screen until subjects clicked on the mouse button to signal that she or he was ready to indicate his or her willingness-to-pay. The subjects could take as long as they wanted to indicate their readiness. Trials where the response time (RT) was 3 standard deviation away from the mean RT of the subject were excluded from further analysis (usually less than 1% of trials), as they could very well indicate disruptions of the experiment outside of the experimenters’ control (e.g., visits from clinicians, nurses, and/or staffs). After making the mouse click, a 1 s fixation period followed. After the fixation period, the subjects would see a price matrix from 0 to 200 in steps of 10 on the computer screen (Fig. 1A). The subjects’ task was to move the cursor to the price box that reflected the most she or he was willing to pay for that food item. Subjects can take as much time as they desired to select one of these boxes with a mouse click, but were encouraged to respond within 2 s. To give the subjects an idea of time, the box where the cursor was at would turn blue within 2 s after the price matrix box appeared. After 2 s, the box where the cursor pointed at would turn red. After clicking on the desired price box, a brief visual feedback on the selected price (willingness-to-pay) was shown (0.5 s), which was then followed by a variable inter-trial interval (1, 1.5, or 2 s).

In this task design, a single trial therefore consisted of two stages: an evaluation stage followed by response stage. During the evaluation stage—when the food item was presented—the subjects were instructed to take time and think over how much money they were willing to pay for the food. By contrast, during the response stage— when the price matrix was shown—the subjects were instructed to indicate willingness-to-pay as quickly as possible. The reason for implementing the two-stage design was to temporally localize the valuation signals we hoped to observe. Since valuation of the food item and the motor response to indicate willingness-to-pay were temporally separated by our task, we hoped that the impact of motor-related confounds introduced when subjects indicated willingness-to-pay could effectively be minimized.

After all trials of the BDM task were complete, one trial was randomly selected and realized according to the rules of the BDM auction. The rules are as follows. Let X be the bid made by a subject for a food item. A random integer Y is drawn from a discrete uniform distribution ranging from 10 to 200 with interval of 10. If X ≥ Y, the subject would buy the food item at a price equal to Y. If X<Y, the subject would not get the food and would keep the endowment. These widely used rules establish a situation whereby the optimal strategy for the subjects is to bid exactly the maximum amount that they are willing to pay for the item. If they underbid for an item, the subject may lose the opportunity to purchase the item later at a still highly desirable price. If they overbid they risk being forced to purchase the item at an undesirable price. Only by stating the exact maximum price at which they would purchase the item can they achieve the optimal result. The BDM rules and the consequences were informed to the subjects before the BDM task so that they knew that the best strategy is to bid exactly what they are willing to pay for the item.

### Electrophysiological recordings

Patients were implanted with 0.86 mm diameter depth electrodes (Ad-Tech, Racine, WI, USA) that were arranged into strips with 6, 8, or 10 contacts (2.29 mm in contact length) and 4-8 mm (most strips: 5 mm) separation. One of the electrode strips was 1.12 mm in diameter with 6 contacts (2.41 mm in contact length) and 5 mm separation in between neighboring contacts.

Recordings were obtained simultaneously from the scalp and depth electrodes while the patients performed the task. Data was collected using the Natus Quantum system (Natus Medical Incorporated). Sampling rates were 2048 Hz with an 878 Hz low-pass filter. During recording, all the electrodes were referenced to the scalp PFz electrode or an intracranial contact located in white matter. Details on the recording sites—MNI coordinates of the electrode contacts—presented in the current study can be found in the Supplementary Tables.

### MRI acquisitions

For each patient, T1-weighed structural MRI images were collected on a 1.5T Signa HDxt scanner (GE Healthcare, Milwaukee, WI, USA) before and after the surgery for electrode implantation. The MR images were taken along the axial plane using a fast spoiled gradient-recalled echo sequence (axial slice thickness=1 mm; TR=10.02 ms, TE=4.28 ms, TI=0 ms, flip angle=15°, bandwidth=31.2 kHz, matrix=256×256, FOV=256×256 mm).

### CT acquisitions

CT images were used in conjunction with T1-weighted MRI images for transforming the anatomical location of the electrode contacts into the standard MNI space. CT images were acquired using Philips Brilliance 64 CT scanner with the following parameters: 64 slices, rotation duration of 1 sec with coverage of 16 cm per rotation, 60-kW generator (512×512 matrix), 120 kV, 301 mAs, and axial slice thickness of 1 mm.

### sEEG data analysis

Below we describe the sEEG data analysis pipeline in detail. A summary of the sEEG data analysis pipeline can be found in the Supplementary Figure 5.

### sEEG preprocessing

EEG data were preprocessed and analyzed by EEGLAB (Delorme & Makeig, 2004) and ERPLAB (Lopez-Calderon & Luck, 2014) in MATLAB in the following steps. First, a digital band-pass filter from 0.5 Hz to 250 Hz and a 60 Hz notch filter were applied to the EEG data at the single contact level— including the scalp and the sEEG dataset—and the EEG data were mean centered. Second, the scalp EEG dataset were separated from the sEEG dataset. Third, the sEEG data for each electrode contact were re-referenced to the average of the two neighboring contacts (Li et al., 2018). The scalp EEG data were re-referenced to the left and right mastoid. Fourth, in order to remove eye-movement-related activity from the sEEG data, we performed an independent component analysis (ICA) separately on scalp EEG dataset and the sEEG dataset. Prior to the ICA, we performed a principal component analysis (PCA) separately to each dataset to denoise the data. Our criterion was to select the number of PCs that explained 95% of the data variance. We then performed ICA on the denoised dataset. Eye-related activities were first identified by inspecting the independent components (ICs) of the scalp EEG data. Once an IC with ocular artifacts was identified, we checked whether there was a corresponding IC in the sEEG data. If there was such a component, it was removed from the sEEG data.

The epoch for each trial (trial epoch) started 1.5 s before the onset of the food stimulus and ended 2 s after stimulus onset, with a pre-stimulus baseline correction. Trial epochs with interictal spikes were identified through visual inspection and were excluded from further data analysis. We referred to the trials with no interictal spikes as the valid trials.

### Time-frequency analysis

After preprocessing, a time frequency analysis was performed using a wavelet transform, estimating spectral power from 4 to 200 Hz for each epoch with full-epoch length single-trial baseline correction (Grandchamp & Delorme, 2011). After time-frequency analysis, the epoch of the timeseries data start at 1 sec before stimulus presentation and end at 1.5 s after stimulus presentation with a 10 ms resolution. The timeseries of the power data from the high-gamma (80-150 Hz), gamma (30-80 Hz), beta (13-30 Hz), alpha (8-12 Hz), theta (4-7 Hz) bands for the epoch were further extracted for the GLM analysis described below (see *General linear modeling of brain activity* below). At each frequency band, the corresponding timeseries data had 251 time points (a 2.5-s time window with a 10ms resolution).

### Identifying the anatomical locations of electrode contacts

To identify the anatomical location of electrode contacts across different subjects, we transformed the electrode contact location from the subject’s native space to standard Montreal Neurological Institute (MNI) space. To do that, we used three sets of brain images collected from each subject: the T1-weighted image prior to electrode implementation (pre-T1), the T1-weighted image after electrode implementation (post-T1), and the CT image after electrode implantation (post-CT). Our goal was to transform the CT image to MNI space. The reason we used the CT image to identify the electrode contact coordinates in the standard space was because the CT image, compared with T1-weighted image, suffers less distortion and therefore allows for more accurate mapping of the contact location. The transformation was performed using SPM12 (Wellcome Trust Centre for Neuroimaging, London, UK; https://www.fil.ion.ucl.ac.uk/spm/) and proceeded in the following three steps. First, the post-CT image was aligned to the post-T1 image with 4^th^ degree B-Spline interpolation. Second, both the post-T1 image and the realigned post-CT image were aligned with the pre-T1 image, also with 4^th^ degree B-Spline interpolation. Finally, the pre-T1, the realigned post-T1 and post-CT images were transformed to the standard MNI space (1 mm isotropic voxel size). The location of each contact in MNI space was obtained through the post-CT images in MNI space.

### Identifying the OFC electrode contacts

We used the Harvard-Oxford probabilistic atlas available in FSL (version 6, https://fsl.fmrib.ox.ac.uk/fsl) to identify the electrode contacts in the OFC. An electrode contact was identified as an OFC contact when the probability of its MNI coordinates being in the OFC was larger than 1%. In addition, because some contacts were situated at the borders between the posterior section of the OFC and the anterior insula, we decided to exclude the contacts that had a higher probability of being in the insula than being in the OFC.

### Visualizing anatomical locations of electrode contacts

To visualize the anatomical location of electrode contacts across different subjects, we used the MNI coordinates of the contacts and plotted them in the standard MNI brain template (Lin et al., 2021; https://github.com/fahsuanlin/fhlin_toolbox).

### General linear modeling of brain activity

For each contact, after time-frequency analysis, we obtained – for each trial – a time-series data of power at a particular frequency band. Here we use high-gamma band (80-150 Hz) as an example and we refer to the power of high-gamma as high-gamma activity, as in Rich and Wallis (2017). To examine subjective-value representations, we performed the following General Linear Modeling (GLM) analysis. First, we set up a GLM for each time point within the epoch separately. Here the data—a vector of length *N_trials_* where *N_trials_* is the number of valid trials a subject performed in the BDM task—are the frequency-specific power obtained from time-frequency analysis (see sEEG preprocessing for descriptions on valid trials). We implemented five different but similar GLMs. The first GLM (GLM-1) was the main GLM, and the rest were slightly different versions of GLM-1 in order to test the robustness of GLM-1’s results on subjective-value representations (Fig. 3). In GLM-1, we implemented a constant term, a regressor for the subjective value of the current trial, and a regressor for the subjective value of the previous trial. In GLM-2, we performed the analysis in two steps. First, we implemented a model with the constant term and the current subjective value regressor. Second, we used the residuals from the first step as data and implemented a model with the constant term and the previous subjective value. In GLM-3, we implemented the same model as GLM-1 except that we only considered trials where the subject’s willingness-to-pay was greater than zero dollars. In GLM-4, the subject’s response time (RT) was added as a regressor, along with the constant term, the current subjective value, and the previous subjective value. In GLM-5, we added the subjective value of the food option encountered two-trials back as a regressor, along with the constant term, the current subjective value, and the previous subjective value. We used the *fitlm* function in MATLAB to perform the GLM analysis.

### Group-level permutation test (across electrode contacts)

This analysis was used for Figs. 2B, 3, 4B, 5B, 6 (middle graphs), and 7 (middle graphs). The data is an *N_contacts_* × *N_time_* matrix of estimated regression coefficients. *N_contacts_* denotes the number of electrode contacts in a region of interest (e.g., orbitofrontal cortex, OFC), and *N_time_* denotes the number of time points within the trial epoch. Using this data, we computed the timeseries of the *t* statistic (across electrode contacts). Hence, the t-statistic timeseries is a 1 × *N_time_* matrix. To compute the *t* statistic, at each time point we computed the mean regression coefficient (across contacts) and divided it by its standard error. We then transformed the *t* statistic to the *z* statistic. To correct for multiple comparisons across time points, we performed a permutation test with threshold-free cluster enhancement (TFCE) (Smith & Nichols, 2009; Winkler et al., 2014). In each permutation, we randomly assigned a label of 1 to half of the contacts and -1 to the other half. Note that for each permutation, this procedure—assigning 1 to half of the contacts and -1 to the other half—was applied to all the time points of the trial epoch. For each time point separately, we then performed a linear regression analysis where data is the actual regression coefficients, and the regressor was the randomly permuted label. This gives us a *t* statistic for the regressor at each time point. The *t* statistic was transformed to the *z* statistic, and using the *z* statistic we computed the TFCE statistic (E=2, H=2) for each time point. As a result, after each permutation, we obtained a timeseries of the TFCE statistic, which we then used to identify the maximum TFCE statistic. After 10,000 permutations, we obtained the null distribution of the maximum TFCE and used it to determine the critical region (p<0.05, familywise error corrected). The TFCE statistic at each time point was then evaluated with respect to the critical region: if the TFCE statistic fell within the critical region, we would conclude that it was statistically significant. Otherwise, it was assessed as not significant.

### Test of symmetry in the data distribution

An important assumption for the permutation test is the symmetry of the data distribution (Winkler et al., 2014). For the group-level permutation tests described above, we had two datasets, one containing the regression coefficients of the current subjective value, and the other for the previous subjective value. Taking the current subjective value as an example, the data is an *N_contacts_* × *N_time_* matrix where each element is the regression coefficient of the current subjective value of a particular electrode contact at a particular point in time in the trial epoch. Each time point consists of a distribution of the regression coefficients across electrode contacts. Our goal was to examine whether this distribution (*N_contacts_* × 1 matrix of regression coefficient) was symmetrical. Therefore, for each time point separately, we computed its corresponding sample skewness (Pearson median skewness). To test symmetry, we used the bootstrap method (resampling with replacement) so as to obtain the distribution of the sample skewness and use it to construct the 95% confidence interval of sample skewness. If the 95% confidence interval did not include 0, we would conclude that the data distribution was symmetrical.

Taking the OFC high-gamma activity as an example: In order to obtain the distribution of the sample skewness, first, we resampled with replacement the OFC contacts (166 in total) 10,000 times. This gave us, for each resampled dataset, a time series of beta coefficients, separately for the current subjective value and the previous subjective value, from the 166 resampled contacts. Second, for each time point separately within the time series, we computed the sample skewness (Pearson median skewness) of the beta coefficients across the resampled OFC contacts. Third, with 10,000 resampled datasets, we obtained, for each time point separately, a distribution of sample skewness that we used to construct the 95% confidence interval. Finally, using the 95% confidence interval, we were able to test whether the data distribution was symmetric at each time point separately. We found that the sample skewness did not differ significantly from 0 in the majority of time points, for each brain region, each frequency band, and for each regressor of interest (current subjective value and previous subjective value). The results are shown in the supplement (Supplementary Figs. 12-14). We did, however, observed a significant positive skewness for the current subjective value in OFC high-gamma activity approximately at 600∼700 ms after stimulus onset (left graph, Supplementary Fig. 12A). On the one hand, this does raise concern for the permutation test regarding the current subjective value at this particular 100-ms time window. On the other hand, we also observed that the time window of activity that significantly correlated with the current subjective value (from approximately 400 ms to 1500 ms after stimulus onset; Fig. 2B) far extended this 100-msec time window. In other words, the significant current subjective-value representations included many time points where the skewness was not significantly different from 0. On this ground, we concluded that the violation of the symmetry assumption observed here should not change the overall conclusion that OFC high-gamma activity represented the current subjective value.

### Group-level permutation test (across contacts) in the time-frequency space

This analysis was used for Fig. 5A. The analysis logic is similar to that described in *Permutation test across contacts in the time domain*. The main difference is the dimensionality of the dataset. Here the dataset included 166 OFC contacts across 20 subjects. In the 1^st^-level GLM, we regressed the power against the current and the previous subjective value for each contact in each time-frequency point. As a result, for the current subjective value and previous subjective value separately, we obtained information about the regression coefficient (which we also referred to as the *beta value*) in a three-dimensional space (time, frequency, electrode contacts). Let N*_freq_* denote the number of frequency points (from 4 Hz to 200 Hz), *N_time_* denotes the number of time points within the trial epoch, and *N_contacts_* denotes the number of electrode OFC contacts. To correct for multiple testing across time points at the group level (across all contacts), we performed permutation test with threshold-free cluster enhancement (TFCE) (Smith & Nichols, 2009; Winkler et al., 2014). In each permutation, we randomly assigned a label of 1 to half of the contacts and -1 to the other half. For each time-frequency point separately, we performed a linear regression analysis (see *General linear modeling of brain activity* above) where data was the actual beta values and the regressor was the randomly permuted label. This would give us the *t* statistic of the regressor at each 2D time-frequency point. The *t* statistic was then transformed to the *z* statistic and based on the z statistic, we computed the TFCE statistic (E=1, H=2) at each 2D time-frequency point. As a result, we obtained a 2D map of TFCE and identified the maximum TFCE. After 10,000 permutations, we obtained the null distribution of the maximum TFCE where we then used to determine whether each 2D time-frequency point was significant (p<0.05, familywise error corrected).

### Individual-level permutation test (for individual electrode contacts)

This analysis was performed for each individual electrode contact separately and was used for Figs. 2C, 4C, 5C, 6 (right graphs), and 7 (right graphs). At each individual contact, the data is a *N_trials_* × *N_time_* matrix of brain activity where *N_trials_* denotes the number of valid trials and *N_time_* the number of time points within the trial epoch. Here brain activity is referred to as the power of a particular frequency band (e.g., high gamma for 80-150 Hz) after time-frequency analysis. For each time point separately, we regressed brain activity (*N_trials_* × 1 matrix) against the current subjective value and the previous subjective value (GLM-1). This gave us a timeseries of regression coefficients for each regressor (1 × *N_time_* matrix) and their corresponding *t* statistics. To correct for multiple testing across time points, we performed a permutation test with threshold-free cluster enhancement (TFCE) (Smith & Nichols, 2009; Winkler et al., 2014). At each permutation, we randomly permuted the trial label (*N_trials_* × 1 matrix) of the design matrix (*N_trials_* × 2 matrix where the current subjective value and previous subjective value were the two regressors). Note that for each permutation, the same permutation was applied to all the time points. We then regressed the data (*N_trials_* × 1 matrix), for each time point separately, against the permuted design matrix. This gave us a *t* statistic of the regressor at each time point. The *t* statistic was transformed to the *z* statistic, and using the *z* statistic we computed the TFCE statistic (E=2, H=2) for each time point. As a result, we obtained a time series of TFCE statistic and identified the maximum TFCE across time. After 10,000 permutations, we obtained the null distribution of the maximum TFCE that we used to determine the critical region (p<0.05, familywise error corrected). If the TFCE statistic of a time point was inside the critical region, it would be labeled as statistically significant. Otherwise, it was labeled as not significant.

### State space analysis

To study how the dynamics of high-gamma activity in the OFC as a whole represented subjective value, we performed two complementary state space analyses, one based on the Principal Component Analysis (PCA) and the other based on a regression subspace analysis (Mante et al., 2013). The data preparation for both analyses was identical and was performed in the following sequence. For each electrode, we first smoothed the high-gamma timeseries data for each trial (Gaussian time window=400 ms). We also performed the same analysis with no smoothing applied, and with 100 ms, 200 ms, and 300 ms Gaussian time window (Supplementary Fig. 9). The timeseries data started from 1s before the onset of the food stimulus and ended 1.5s after stimulus onset. Second, we computed the average timeseries (across all trials) and subtracted it from the timeseries of each trial. Third, we sorted the trials into four conditions according to the magnitude of the subjective value of the current trial (high or low, median spit) and the magnitude of the subjective value on the previous trial (high or low, median spilt). The medians were obtained based on the corresponding subject’s willingness-to-pay data. The four conditions therefore are [current subjective value, previous subjective value] = [high, low], [high, high], [low, low], [low, high]. Fourth, we computed the average timeseries of each condition. Fifth, for each condition, we organized the average timeseries data of all electrode contacts as a two-dimensional matrix of size *N_contacts_* × *N_time_* where *N_contacts_* is the number of electrode contacts in the OFC across all subjects (*N_contacts_* = 166) and *N_time_* is the number of time points within the trial epoch (*N_time_* = 251). We denote this condition-specific matrix *X_c_*. Finally, we collapsed the four 2D activity matrices (one from each condition) such that the final dataset for the subsequent analyses (PCA-based analysis and regression subspace analysis described below) was a 2D matrix of size 166 × 1044 which included all data aggregated together before the initial PCA was performed. We denote this data matrix *X*.

#### PCA-based analysis

We performed PCA on the prepared dataset described above using the *pca* function in MATLAB. The feature dimensions were the electrode contacts, and the observations were the time points within the trial epoch. We then projected the activity matrix of each condition onto the first two PCs, resulting in four different trajectories (timeseries) in the PC space. We then plotted the trajectories and color coded them (Fig. 8B-D).

#### Regression subspace analysis

We first performed PCA on the prepared dataset to denoise the data. In the main text, the prepared dataset was the smoothed high-gamma timeseries data (Gaussian time window of 400 ms). We also performed the analysis with no smoothing applied (Supplementary Fig. 11). The number of PCs selected to construct the denoising matrix was 12. We also performed the same analysis described below with number of PCs being 2 and 20 (Supplementary Fig. 10). The denoising matrix (*D*) was constructed

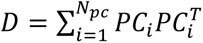

where *PC_i_* is the i-th principal component and is a column vector of size *N_contacts_*. The resulting *D* is a *N_contacts_* × *N_contacts_* matrix. The denoised data *X_pca_* is obtained according to

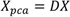

Next, we turn our attention to the linear regression analysis (GLM-1) and the regression coefficients. In GLM-1, we implemented two task-related regressors, namely the subjective value of the current trial and the subjective value of the previous trial. The GLM was performed on each electrode contact and for each time point within the trial epoch separately. Let *v* denote task-related variable. Here we have two task related variables, the current and the previous subjective value. Let *β_v,t_* denote the regression vector consisting of regression coefficient of each contact associated with task variable *v* (current subjective value or previous subjective value) at time *t*. *β*_*v,t*_ therefore has a length of *N_contacts_*. We then applied the denoising matrix (*D*) to *β*_*v,t*_ to denoise the regression vector

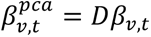

where 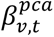 is the denoised regression vector.

For each task variable *v*, we find the time 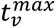 that has the maximum L2 norm of 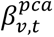 and define the corresponding regression vector 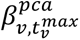 as 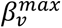 where 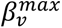 is the time-independent, de-noised regressor vector for task variable *v*. We then put *β*^*%5^ from different task variables (current and previous subjective value) together into a single 2D matrix *β*^max^ where the columns are the task variables and the rows are the electrode contacts. *β*^max^ therefore has a size of *N_v_* × *N_contacts_* where *N_v_* = 2 and *N_contacts_* = 166. Finally, we obtain the orthogonal axes of the current subjective value and the previous subjective value by orthogonalizing *β^max^* with QR decomposition

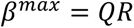

where *Q* is an orthogonal matrix and *R* is an upper triangular matrix. The first two columns of Q correspond to the orthogonalized regressor vectors 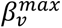, which we refer to as the task-related axes of the current subjective value and the previous subjective value.

To study the representations of the current and previous subjective value in the OFC, we projected the condition-specific data matrix *Xc* onto the orthogonal axes

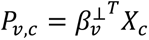

Where *P_v,c_* is the set of timeseries vectors over all task variables (*v*) and conditions (*c*). Here we have two task variables, the current and previous subjective value, and four conditions. Therefore, in the two-dimensional space with the current and previous subjective value as the task-related axes, we have four trajectories, which we plotted in Fig. 8E-G.

## Supplementary Material

**Supplementary Figure 1.**
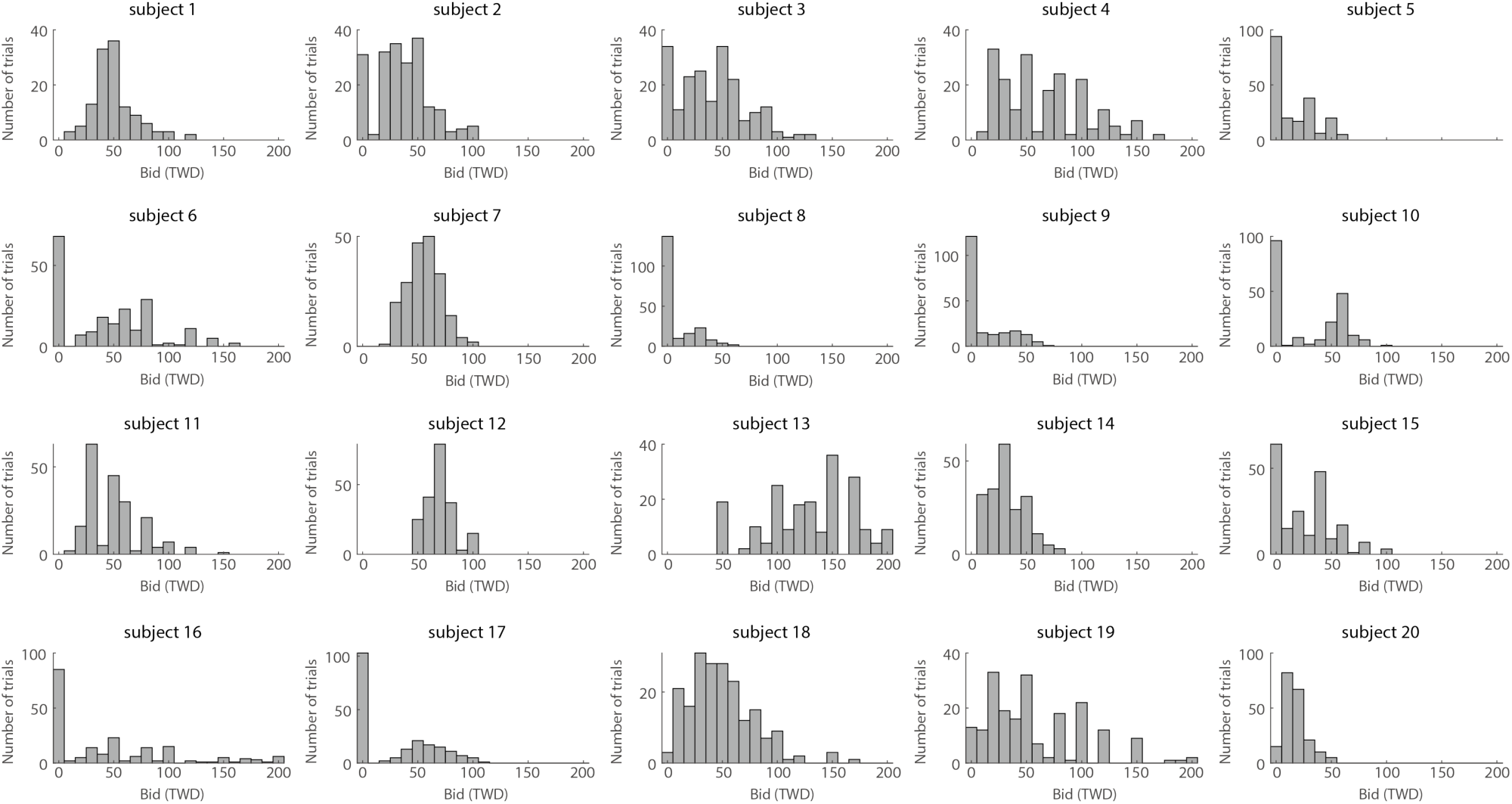
The distribution of willingness-to-pay for individual subjects.

**Supplementary Figure 2.**
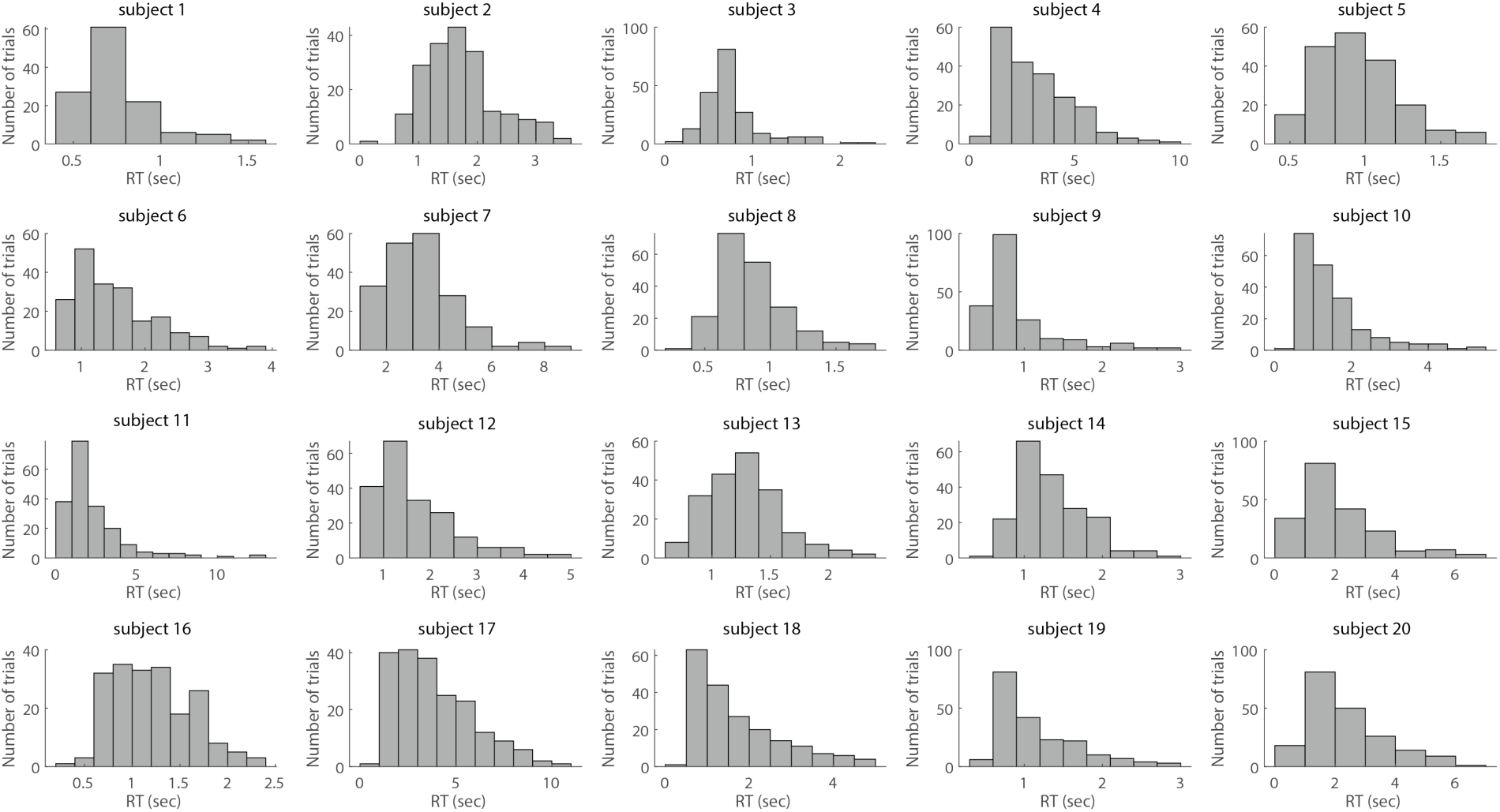
The response time (RT) distribution of individual subjects.

**Supplementary Figure 3.**
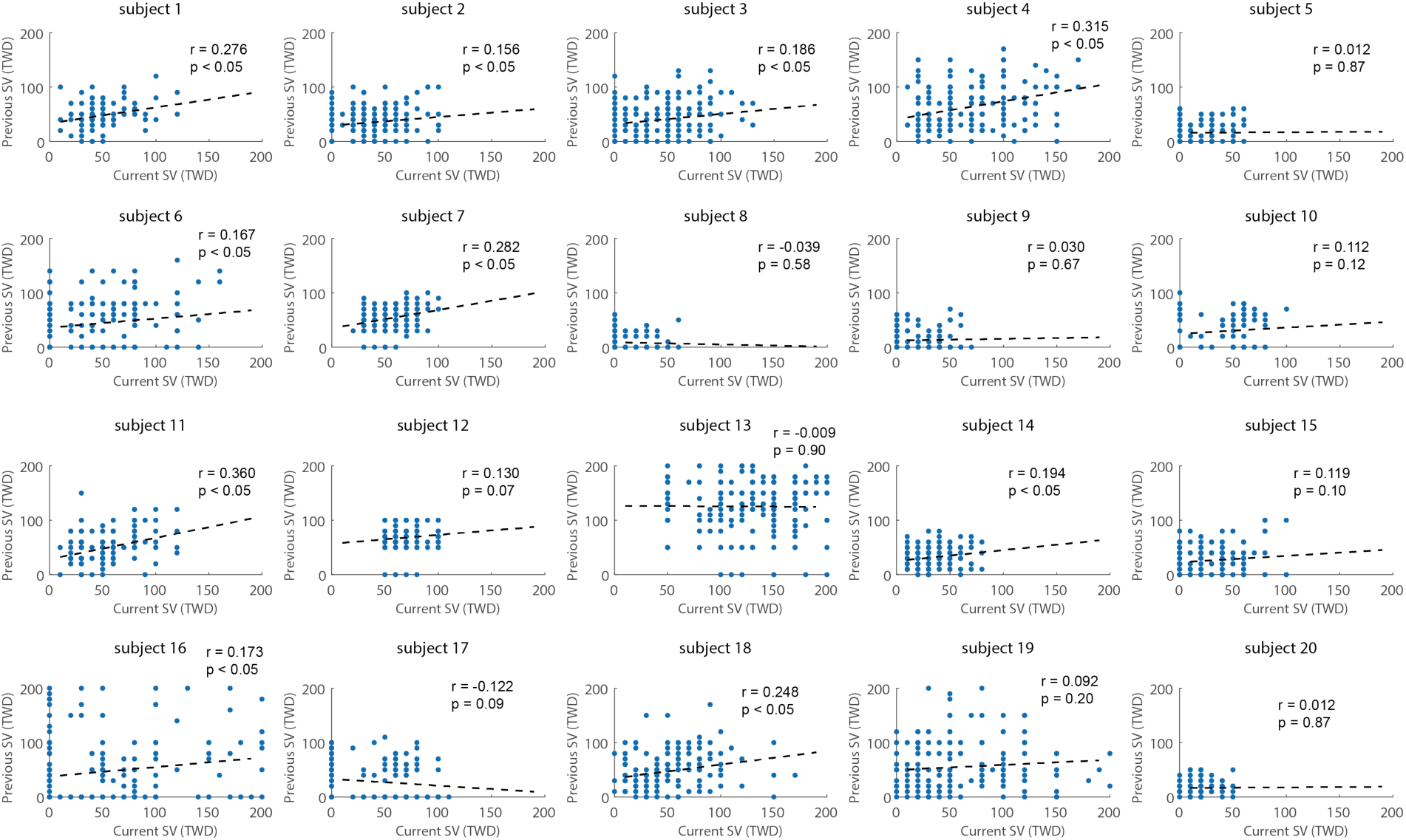
Correlation between the subjective value of the current trial and the subjective value of the previous trial. *r* indicates the Pearson correlation coefficient, and *p* indicates the corresponding *p*-value.

**Supplementary Figure 4.**
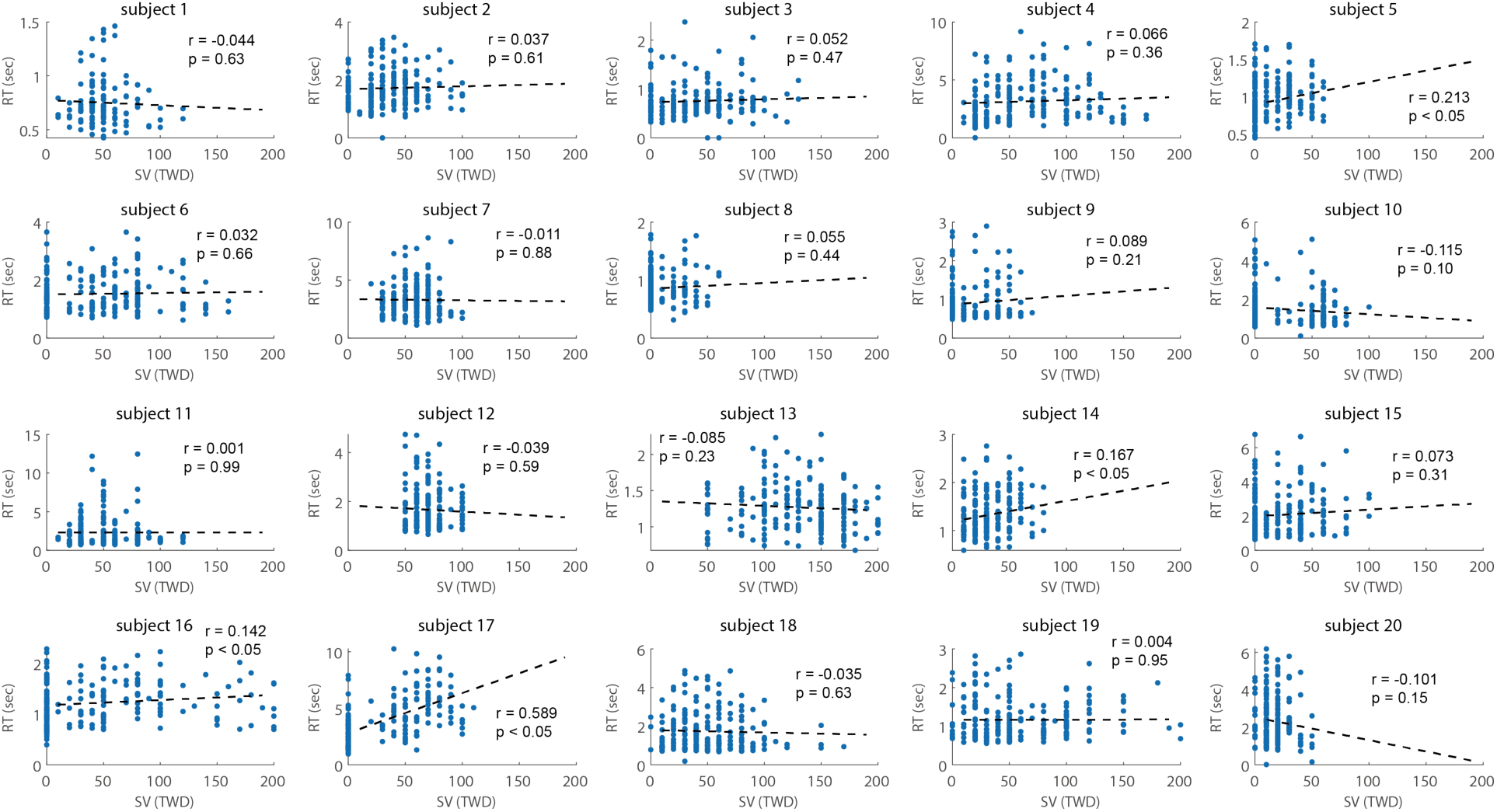
Correlation between the subjective value of the current trial and the response time (RT). *r* indicates the Pearson correlation coefficient, and *p* indicates the *p*-value.

**Supplementary Figure 5.**
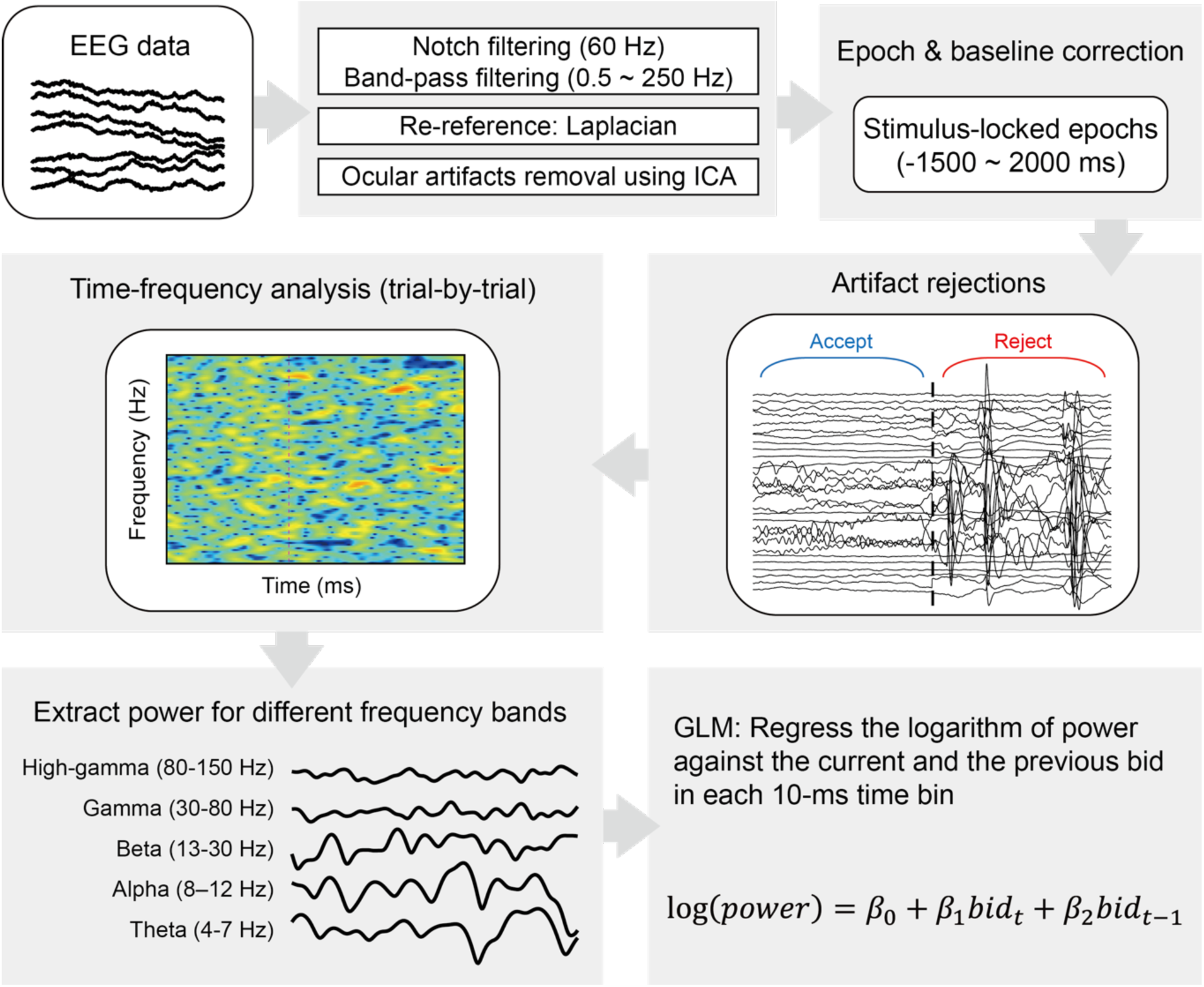
Overview of the sEEG data processing pipeline. First, the sEEG data were filtered to remove uninteresting frequency bands and power-line interference. Second, the sEEG data of each electrode contact were re-referenced to the average of the two neighboring contacts (Laplacian reference). Third, the ocular artifacts were identified and removed from the sEEG data using the Independent Component Analysis (ICA). The trial epoch for each trial was 3.5 s, from 1.5 s before the onset of stimulus to 2 s after stimulus onset. The baseline used for baseline correction was the 2-s pre-stimulus interval. The epochs with interictal activities were identified and excluded from further analysis. After preprocessing, a time-frequency analysis was performed for each epoch. The time series of the power from different frequency bands were then extracted. For each contact, we regressed the logarithm of the power in each 10 ms time bin against the regressors – the bid (willingness-to-pay) subjects revealed in the current trial (*bidt*) and that in the previous trial (*bidt-1*). The estimated regression coefficients (*β*_1_, *β*_2_) reflect the strength and direction of the representations for the subjective value of the current trial (*β*_1_) and that of the previous trial (*β*_2_).

**Supplementary Figure 6.**
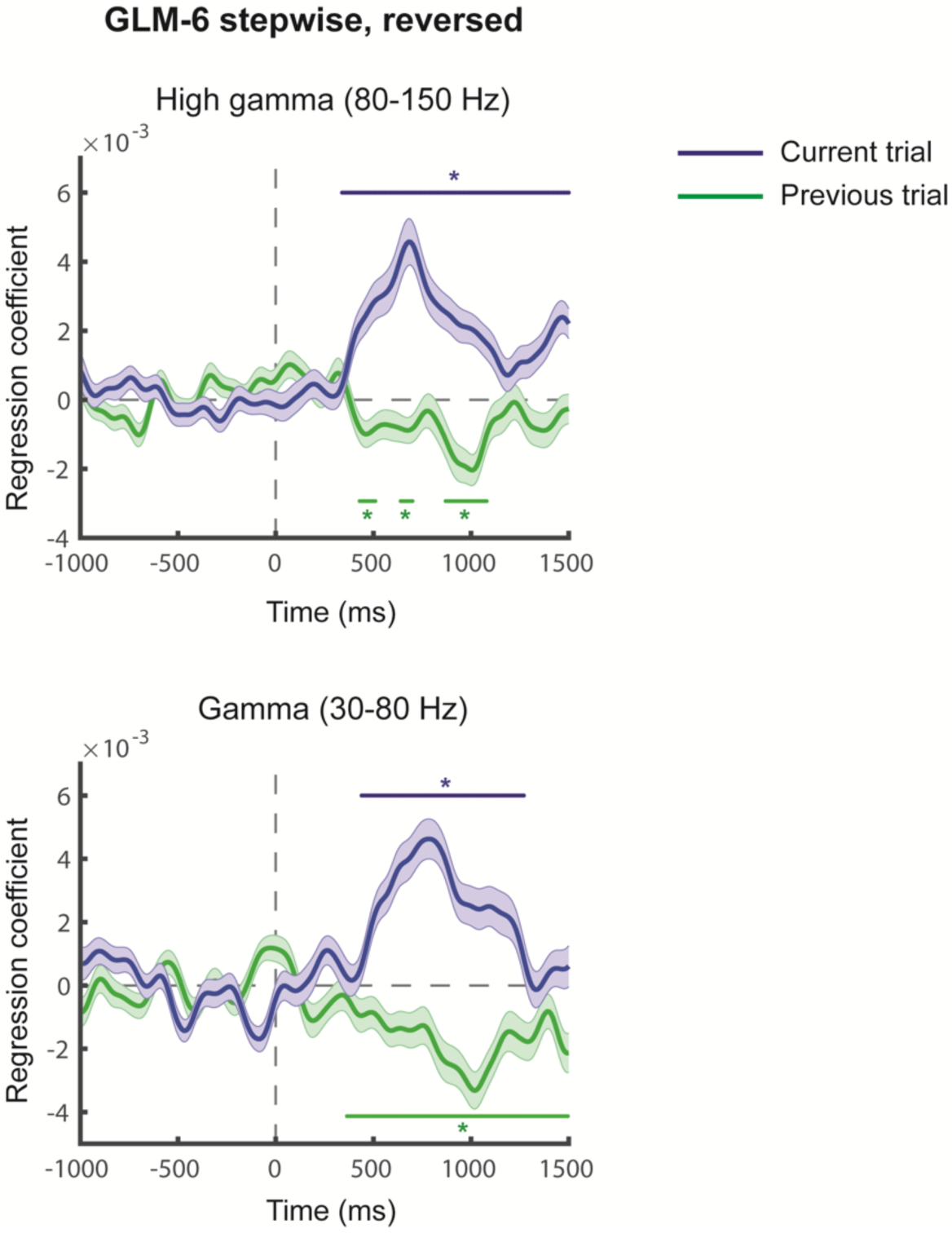
Testing the robustness of subjective-value representations in the OFC. Here we revered the order of steps in GLM-2. In the first step, we regressed high-gamma activity against the previous subjective value. In the second step, we used the residuals from the first step as data and regressed them against the current subjective value. Results were similar to GLM-2 (Fig. 3 in the main text). Conventions are the same as described in Fig. 3 in the main text.

**Supplementary Figure 7.**
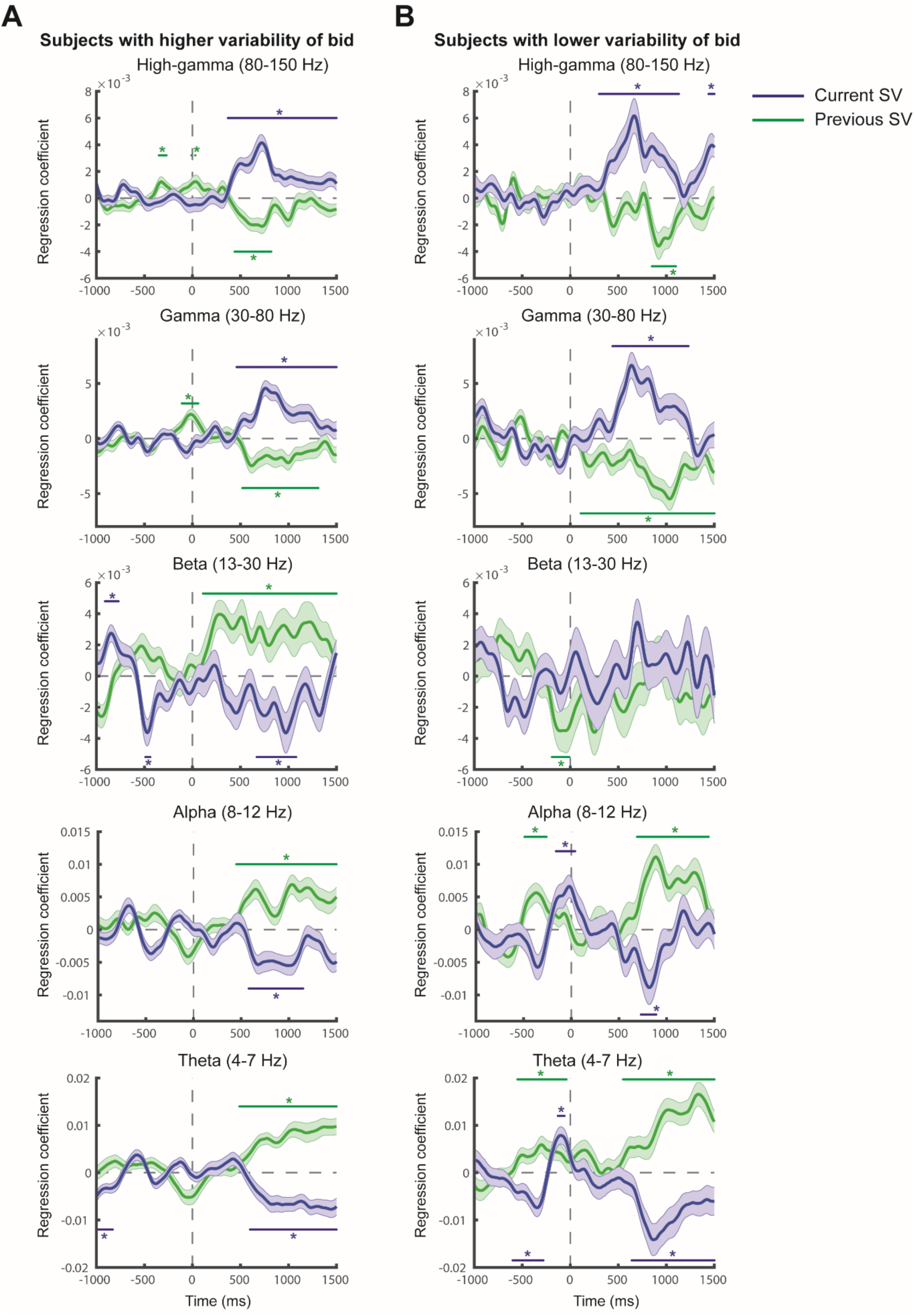
Subjective-value representations in the OFC for subjects with different variability of bids. We split the subjects to two different groups according to the variability of their bids (willingness-to-pay). A. The 10 subjects with larger variability of bids (94 contacts). B. The 10 subjects with lower variability of bids (72 contacts). Conventions are the same as in Figs. 3 and 5 in the main text.

**Supplementary Figure 8.**
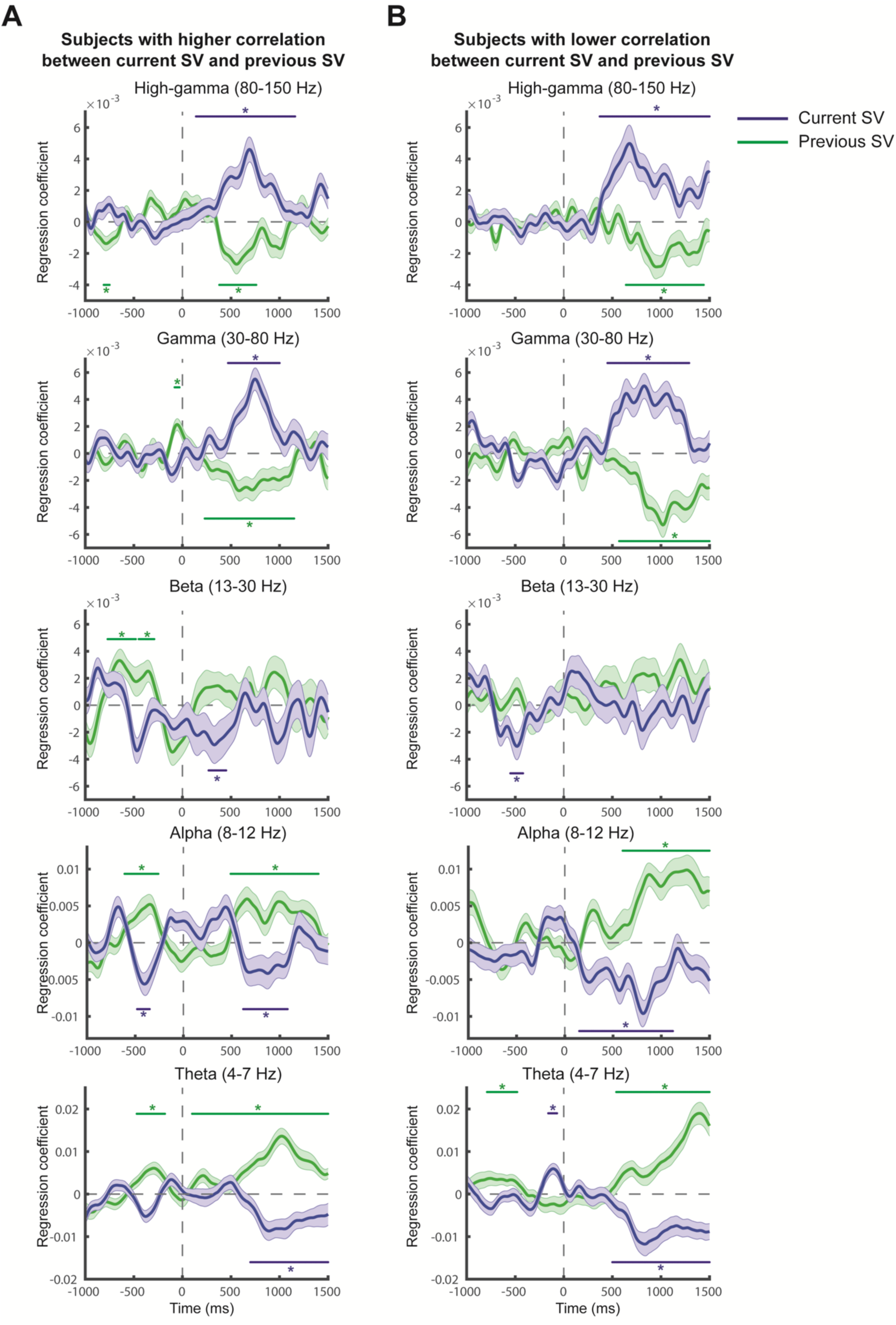
Subjective-value representations in the OFC for subjects with different degree of correlations between the current subjective value and the previous subjective value. We split the subjects into two groups according to the degree of correlation. A. The 10 subjects with larger correlation between the current subjective value and the previous subjective value (87 contacts). B. The 10 subjects with lower correlation between current subjective value and the previous subjective value (79 contacts). Conventions are the same as described in Figs. 3 and 5 in the main text.

**Supplementary Figure 9.**
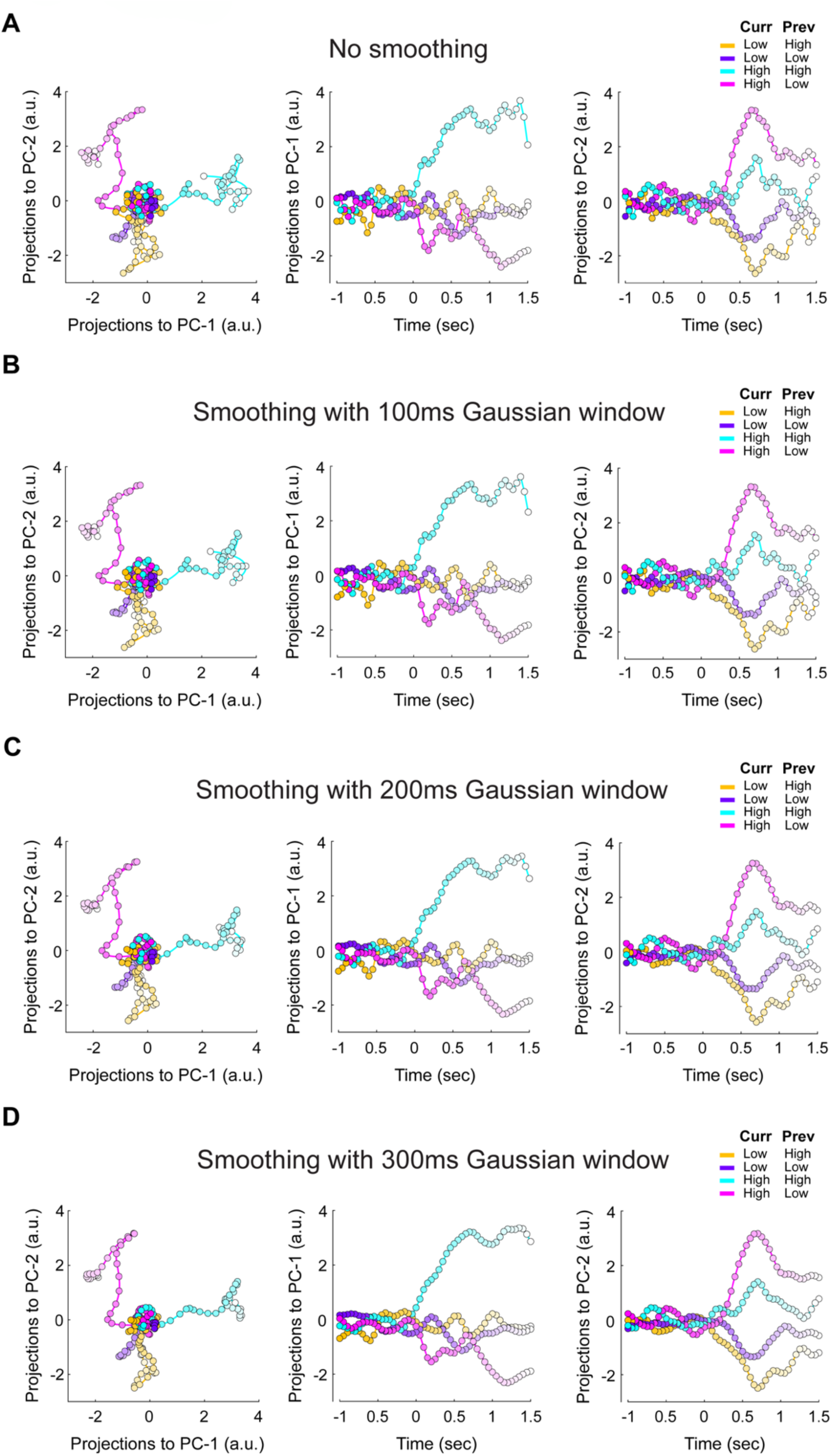
Trajectories of OFC population high-gamma activity in the space of the first and second principal components (PC-1, PC-2) under different smoothing parameters. **A.** No smoothing was applied to the trial-level high-gamma timeseries data. **B.** Data were smoothed with a 100ms Gaussian window. **C.** Data were smoothed with a 200ms Gaussian window. **D.** Data were smoothed with a 300ms Gaussian window.

**Supplementary Figure 10.**
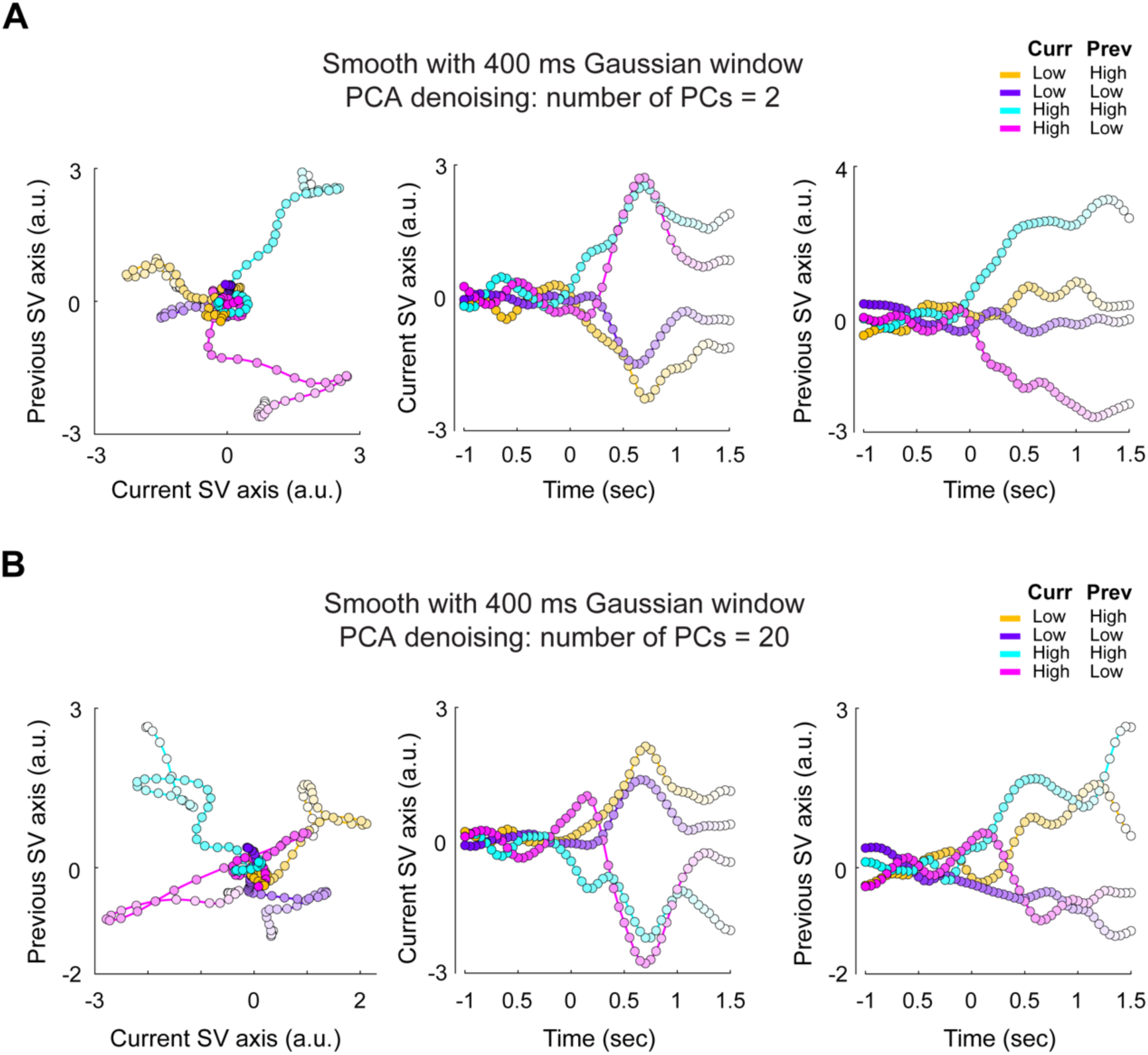
Regression subspace analysis. Trajectories of OFC population high-gamma activity in the space of current subjective value (Current SV axis) and previous subjective value (Previous SV axis) under different PCA-based denoising setup when the trial-level high-gamma timeseries data were smoothed with a 400ms Gaussian time window. A. When the number of PCs included is 2. B. When the number of PCs included is 20.

**Supplementary Figure 11.**
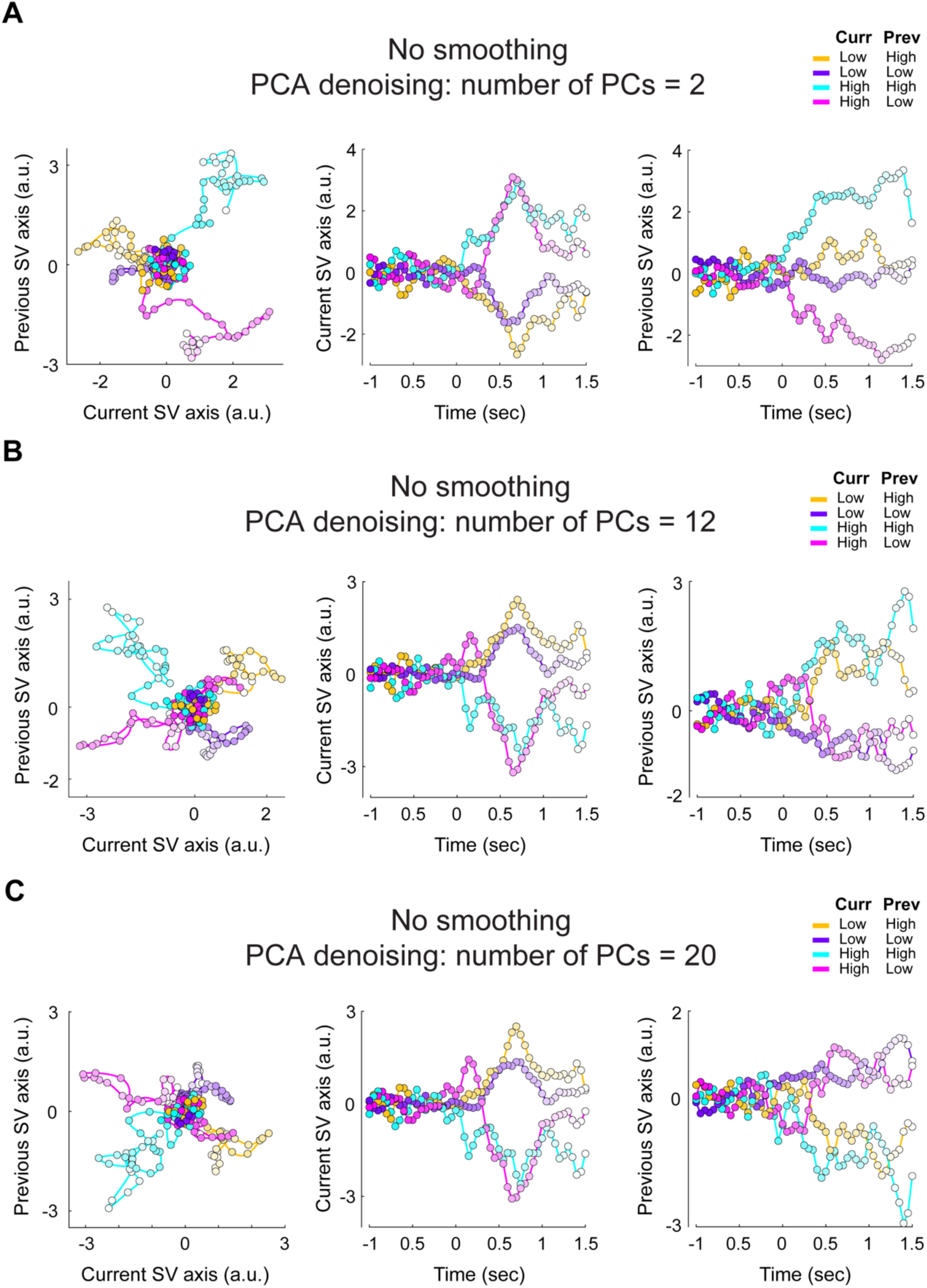
Regression subspace analysis. Trajectories of OFC population high-gamma activity in the space of current subjective value (Current SV axis) and previous subjective value (Previous SV axis) under different PCA-based denoising setup when no smoothing was applied to the trial-level high-gamma timeseries data. A. The number of PCs included in the denoising matrix is 2. B. 12 PCs were included in the denoising matrix. C. 20 PCs were included in the denoising matrix.

**Supplementary Figure 12.**
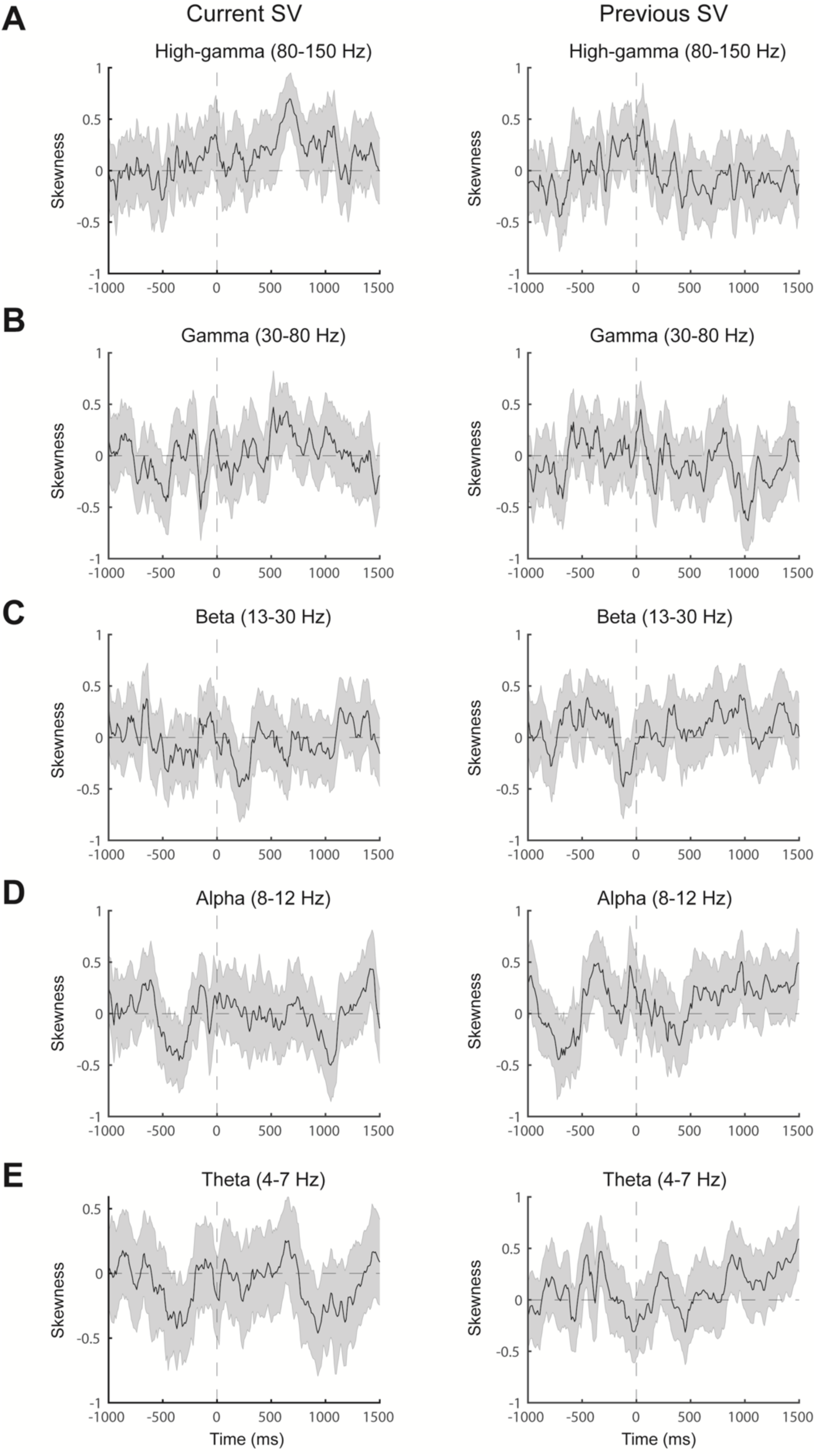
Sample skewness of the regression coefficient in GLM-1 in the OFC contacts. **A.** High-gamma power (80-150 Hz). **B.** Gamma power (30-80 Hz). **C.** Beta power (13-30 Hz). D. Alpha power (8-12 Hz). E. Theta power (4-7 Hz).

**Supplementary Figure 13.**
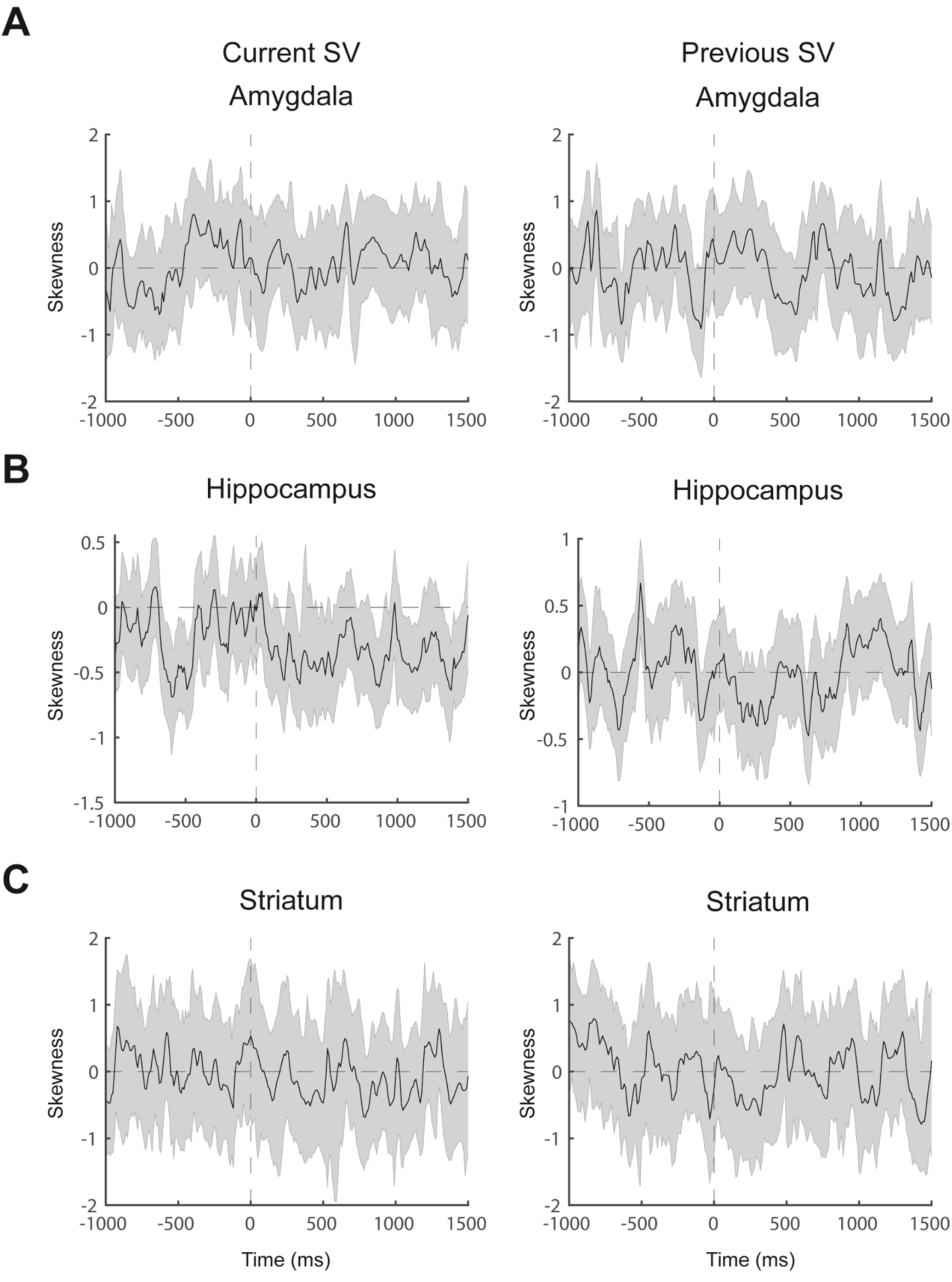
Sample skewness of the regression coefficient in GLM-1 in the subcortical contacts. **A.** Amygdala (30 contacts). **B.** Hippocampus (126 contacts). **C.** Striatum (25 contacts).

**Supplementary Figure 14.**
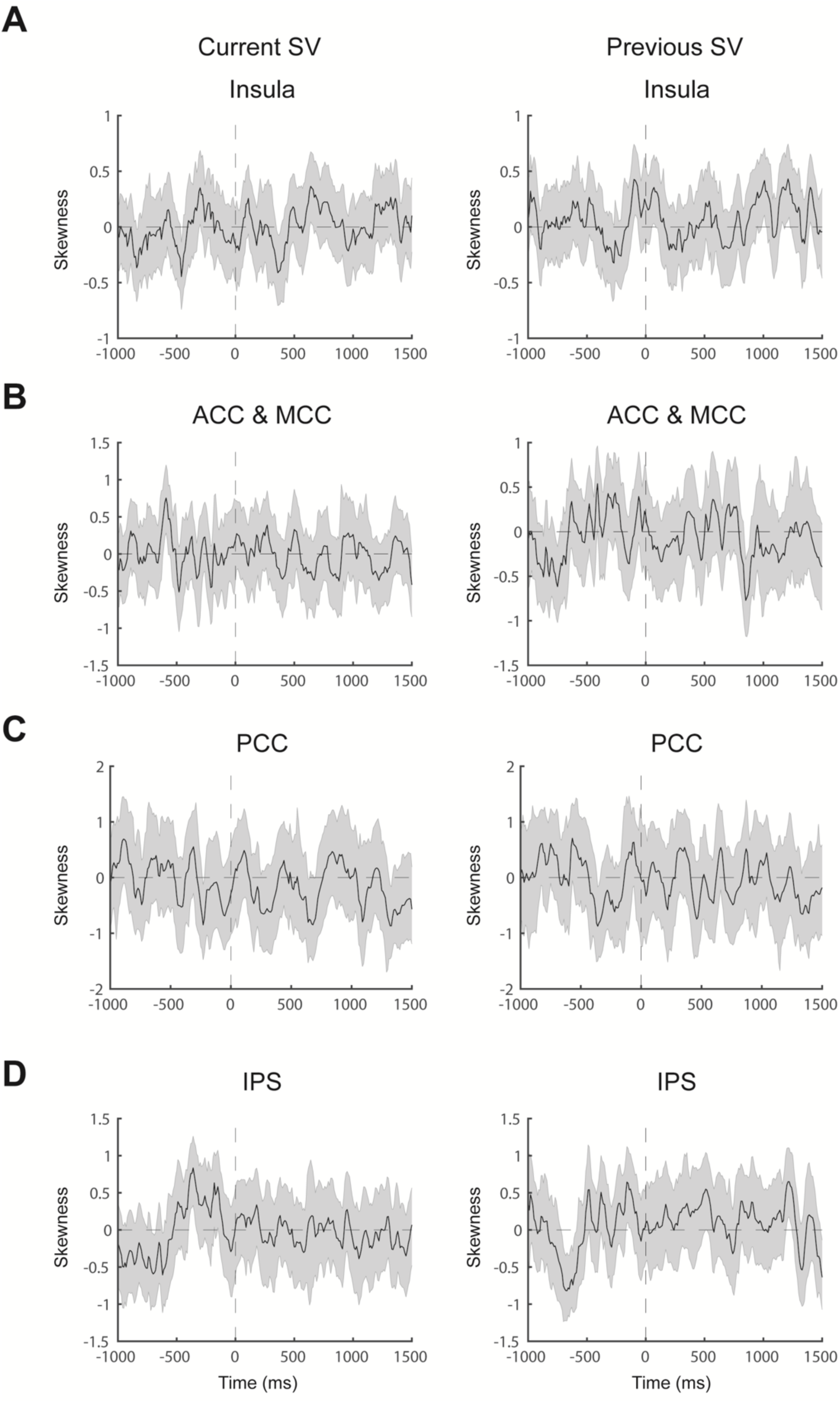
Sample skewness of the regression coefficient in GLM-1 in the cortical contacts. A. Insula (169 contacts). B. Anterior cingulate and midcingulate cortex (ACC and MCC, 81 contacts). C. Posterior cingulate cortex (PCC, 31 contacts). D. Intraparietal sulcus (IPS, 62 contacts).

**Supplementary Table 1:**
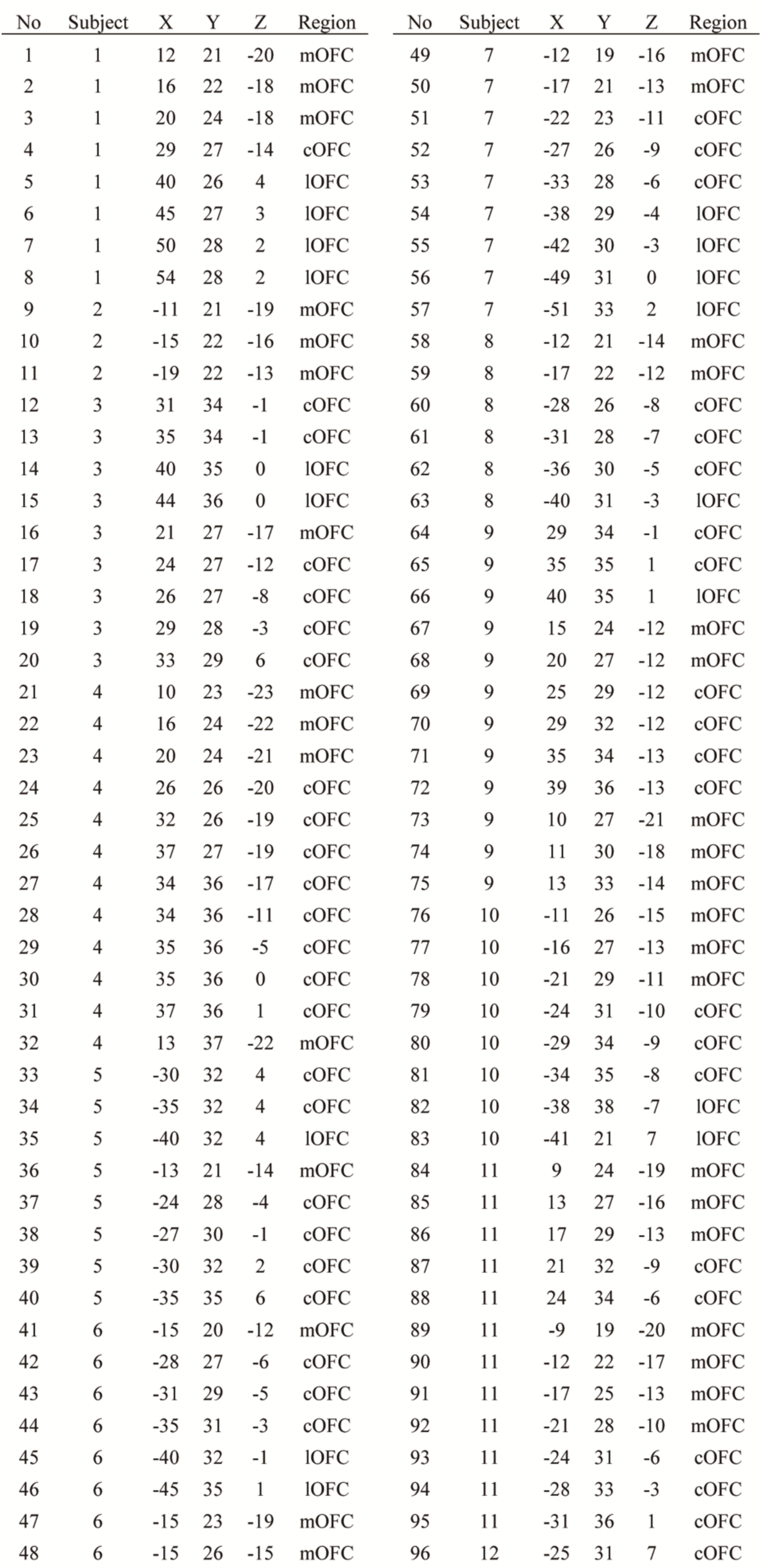
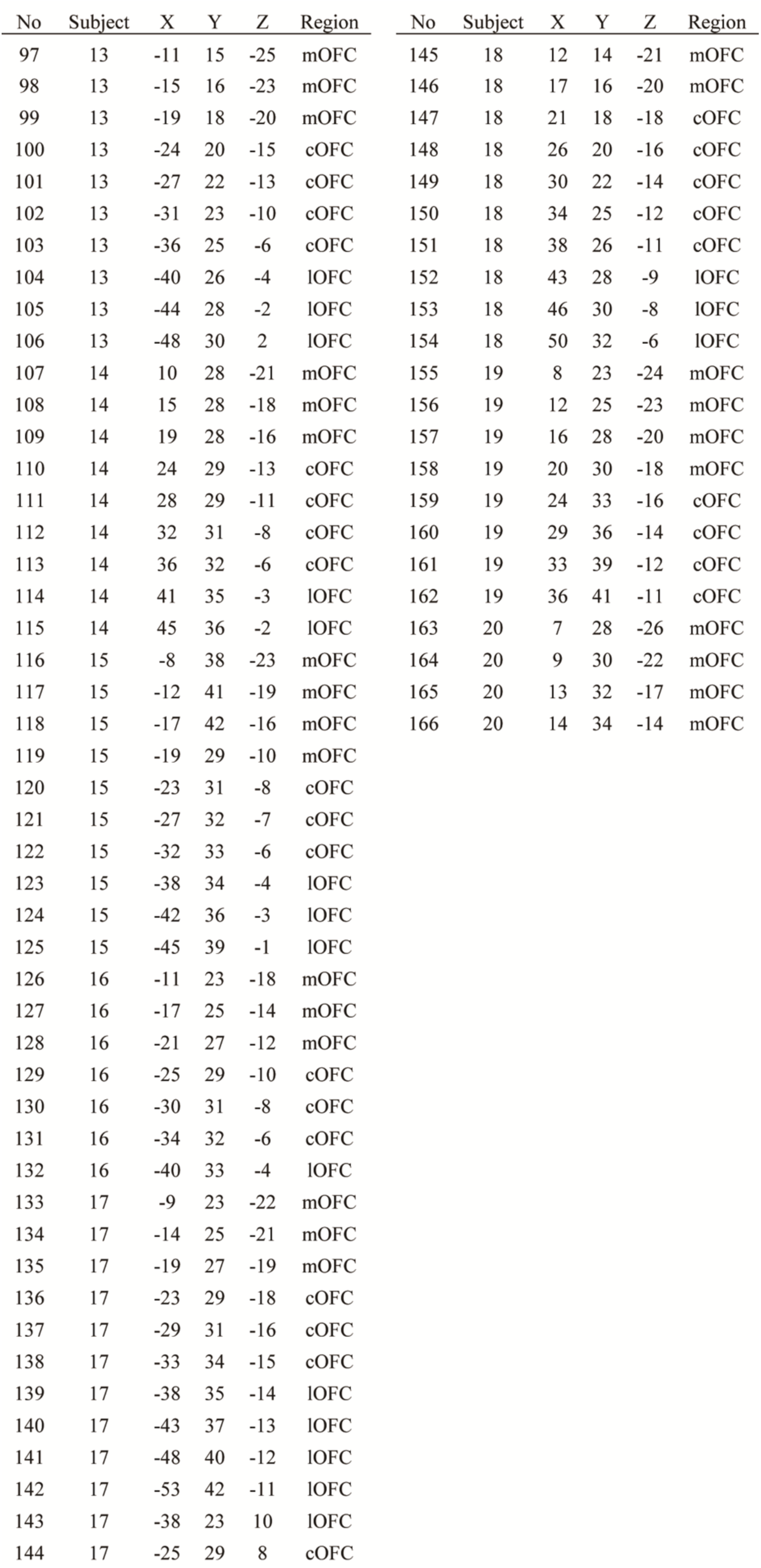
Orbitofrontal cortex (OFC). MNI coordinates for the 166 OFC contacts.

**Supplementary Table 2:** Amygdala. MNI coordinates for the 30 amygdala contacts.

| No | Subject | X | Y | Z |
| --- | --- | --- | --- | --- |
| 1 | 1 | 23 | -2 | -22 |
| 2 | 1 | 27 | 0 | -22 |
| 3 | 1 | 32 | 2 | -23 |
| 4 | 2 | -25 | -4 | -21 |
| 5 | 2 | -29 | -3 | -20 |
| 6 | 2 | -34 | -2 | -20 |
| 7 | 4 | 23 | -4 | -27 |
| 8 | 4 | 28 | -2 | -26 |
| 9 | 4 | 33 | -1 | -26 |
| 10 | 5 | -27 | -6 | -23 |
| 11 | 5 | -32 | -4 | -22 |
| 12 | 11 | 26 | -4 | -23 |
| 13 | 11 | 31 | -3 | -23 |
| 14 | 11 | -21 | -5 | -24 |
| 15 | 11 | -27 | -4 | -24 |
| 16 | 11 | -31 | -4 | -24 |
| 17 | 13 | -35 | 0 | -18 |
| 18 | 13 | -28 | -4 | -25 |
| 19 | 13 | -33 | -3 | -25 |
| 20 | 15 | -33 | -6 | -25 |
| 21 | 16 | -30 | -4 | -25 |
| 22 | 17 | -18 | -5 | -24 |
| 23 | 17 | -23 | -4 | -25 |
| 24 | 17 | -29 | -3 | -25 |
| 25 | 19 | 17 | -1 | -29 |
| 26 | 19 | 22 | -1 | -28 |
| 27 | 19 | 26 | 0 | -27 |
| 28 | 19 | 30 | 1 | -27 |
| 29 | 20 | 29 | -5 | -23 |
| 30 | 20 | 34 | -4 | -23 |

**Supplementary Table 3:** **Hippocampus.** MNI coordinates for 126 hippocampus contacts.

| No | Subject | X | Y | Z | No | Subject | X | Y | Z | No | Subject | X | Y | Z |
| --- | --- | --- | --- | --- | --- | --- | --- | --- | --- | --- | --- | --- | --- | --- |
| 1 | 1 | 25 | -16 | -20 | 49 | 10 | -32 | -15 | -20 | 97 | 16 | -27 | -31 | -14 |
| 2 | 1 | 30 | -15 | -20 | 50 | 10 | -38 | -15 | -19 | 98 | 16 | -33 | -32 | -13 |
| 3 | 1 | 35 | -14 | -20 | 51 | 11 | 21 | -7 | -22 | 99 | 16 | -37 | -31 | -13 |
| 4 | 1 | 39 | -13 | -20 | 52 | 11 | 26 | -17 | -22 | 100 | 17 | -34 | -2 | -25 |
| 5 | 1 | 22 | -28 | -14 | 53 | 11 | 29 | -16 | -21 | 101 | 18 | 23 | -7 | -29 |
| 6 | 1 | 31 | -28 | -14 | 54 | 11 | 35 | -15 | -18 | 102 | 18 | 28 | -6 | -29 |
| 7 | 1 | 36 | -28 | -15 | 55 | 11 | 23 | -37 | -7 | 103 | 18 | 33 | -5 | -29 |
| 8 | 2 | -32 | -18 | -20 | 56 | 11 | 28 | -37 | -7 | 104 | 18 | 26 | -17 | -21 |
| 9 | 2 | -37 | -17 | -19 | 57 | 11 | 33 | -36 | -7 | 105 | 18 | 31 | -17 | -21 |
| 10 | 2 | -30 | -32 | -7 | 58 | 11 | 38 | -35 | -6 | 106 | 18 | 35 | -16 | -21 |
| 11 | 2 | -35 | -33 | -8 | 59 | 11 | -23 | -18 | -18 | 107 | 18 | 40 | -15 | -21 |
| 12 | 4 | 27 | -15 | -18 | 60 | 11 | -28 | -18 | -18 | 108 | 18 | 31 | -33 | -15 |
| 13 | 4 | 32 | -14 | -17 | 61 | 11 | -33 | -19 | -18 | 109 | 18 | 35 | -33 | -13 |
| 14 | 4 | 38 | -14 | -17 | 62 | 11 | -20 | -33 | -11 | 110 | 19 | 21 | -12 | -27 |
| 15 | 4 | 23 | -31 | -9 | 63 | 11 | -24 | -34 | -11 | 111 | 19 | 25 | -12 | -27 |
| 16 | 4 | 28 | -33 | -9 | 64 | 11 | -28 | -36 | -10 | 112 | 19 | 30 | -12 | -25 |
| 17 | 4 | 32 | -34 | -8 | 65 | 11 | -34 | -37 | -9 | 113 | 19 | 35 | -12 | -24 |
| 18 | 4 | 38 | -34 | -7 | 66 | 12 | -24 | -11 | -20 | 114 | 19 | 24 | -28 | -17 |
| 19 | 5 | -18 | -9 | -24 | 67 | 12 | -28 | -11 | -20 | 115 | 19 | 29 | -28 | -17 |
| 20 | 5 | -23 | -8 | -23 | 68 | 12 | -33 | -10 | -21 | 116 | 19 | 34 | -28 | -18 |
| 21 | 5 | -21 | -17 | -21 | 69 | 12 | -23 | -20 | -17 | 117 | 20 | 18 | -6 | -24 |
| 22 | 5 | -26 | -17 | -21 | 70 | 12 | -27 | -20 | -18 | 118 | 20 | 24 | -6 | -23 |
| 23 | 5 | -31 | -17 | -21 | 71 | 12 | -31 | -20 | -17 | 119 | 20 | 24 | -17 | -20 |
| 24 | 5 | -35 | -16 | -21 | 72 | 12 | -36 | -20 | -16 | 120 | 20 | 28 | -17 | -20 |
| 25 | 5 | -26 | -30 | -6 | 73 | 12 | -20 | -30 | -16 | 121 | 20 | 33 | -16 | -20 |
| 26 | 5 | -30 | -31 | -6 | 74 | 12 | -25 | -30 | -15 | 122 | 20 | 38 | -16 | -19 |
| 27 | 5 | -35 | -32 | -6 | 75 | 12 | -29 | -30 | -14 | 123 | 20 | 23 | -24 | -16 |
| 28 | 6 | -20 | -12 | -24 | 76 | 12 | -33 | -31 | -12 | 124 | 20 | 28 | -24 | -15 |
| 29 | 6 | -25 | -12 | -24 | 77 | 12 | -37 | -30 | -12 | 125 | 20 | 33 | -23 | -14 |
| 30 | 6 | -30 | -12 | -24 | 78 | 12 | 20 | -17 | -21 | 126 | 20 | 38 | -24 | -14 |
| 31 | 6 | -35 | -14 | -23 | 79 | 12 | 25 | -15 | -21 |  |  |  |  |  |
| 32 | 6 | -23 | -27 | -16 | 80 | 12 | 29 | -13 | -21 |  |  |  |  |  |
| 33 | 6 | -28 | -27 | -15 | 81 | 12 | 33 | -12 | -22 |  |  |  |  |  |
| 34 | 6 | -33 | -26 | -14 | 82 | 12 | 37 | -10 | -22 |  |  |  |  |  |
| 35 | 6 | -38 | -27 | -13 | 83 | 12 | 30 | -24 | -23 |  |  |  |  |  |
| 36 | 7 | -26 | -38 | -13 | 84 | 13 | -35 | -18 | -23 |  |  |  |  |  |
| 37 | 8 | -19 | -10 | -23 | 85 | 15 | -23 | -7 | -25 |  |  |  |  |  |
| 38 | 8 | -24 | -9 | -23 | 86 | 15 | -28 | -7 | -25 |  |  |  |  |  |
| 39 | 8 | -28 | -8 | -24 | 87 | 15 | -22 | -18 | -18 |  |  |  |  |  |
| 40 | 8 | -33 | -8 | -25 | 88 | 15 | -26 | -18 | -17 |  |  |  |  |  |
| 41 | 8 | -22 | -17 | -19 | 89 | 15 | -31 | -18 | -17 |  |  |  |  |  |
| 42 | 8 | -27 | -18 | -18 | 90 | 15 | -36 | -19 | -17 |  |  |  |  |  |
| 43 | 8 | -31 | -20 | -18 | 91 | 16 | -20 | -7 | -26 |  |  |  |  |  |
| 44 | 8 | -24 | -35 | -14 | 92 | 16 | -25 | -6 | -26 |  |  |  |  |  |
| 45 | 8 | -29 | -35 | -13 | 93 | 16 | -24 | -18 | -21 |  |  |  |  |  |
| 46 | 8 | -34 | -35 | -12 | 94 | 16 | -29 | -18 | -20 |  |  |  |  |  |
| 47 | 10 | -23 | -15 | -22 | 95 | 16 | -34 | -17 | -20 |  |  |  |  |  |
| 48 | 10 | -29 | -15 | -20 | 96 | 16 | -23 | -30 | -15 |  |  |  |  |  |

**Supplementary Table 4:**
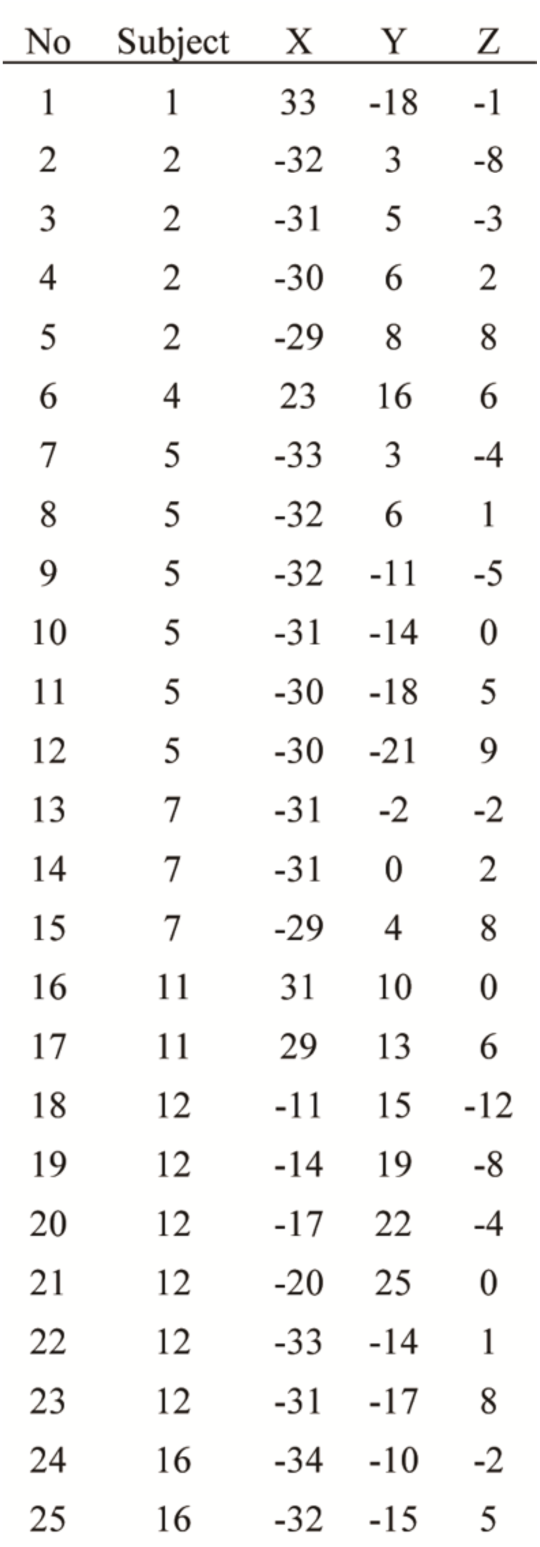
Striatum. MNI coordinates for the 25 striatum contacts.

| No | Subject | X | Y | Z |
| --- | --- | --- | --- | --- |
| 1 | 1 | 33 | -18 | -1 |
| 2 | 2 | -32 | 3 | -8 |
| 3 | 2 | -31 | 5 | -3 |
| 4 | 2 | -30 | 6 | 2 |
| 5 | 2 | -29 | 8 | 8 |
| 6 | 4 | 23 | 16 | 6 |
| 7 | 5 | -33 | 3 | -4 |
| 8 | 5 | -32 | 6 | 1 |
| 9 | 5 | -32 | -11 | -5 |
| 10 | 5 | -31 | -14 | 0 |
| 11 | 5 | -30 | -18 | 5 |
| 12 | 5 | -30 | -21 | 9 |
| 13 | 7 | -31 | -2 | -2 |
| 14 | 7 | -31 | 0 | 2 |
| 15 | 7 | -29 | 4 | 8 |
| 16 | 11 | 31 | 10 | 0 |
| 17 | 11 | 29 | 13 | 6 |
| 18 | 12 | -11 | 15 | -12 |
| 19 | 12 | -14 | 19 | -8 |
| 20 | 12 | -17 | 22 | -4 |
| 21 | 12 | -20 | 25 | 0 |
| 22 | 12 | -33 | -14 | 1 |
| 23 | 12 | -31 | -17 | 8 |
| 24 | 16 | -34 | -10 | -2 |
| 25 | 16 | -32 | -15 | 5 |

**Supplementary Table 5:** Insula. MNI coordinates for the 169 insula contacts.

| No | Subject | X | Y | Z | No | Subject | X | Y | Z | No | Subject | X | Y | Z |
| --- | --- | --- | --- | --- | --- | --- | --- | --- | --- | --- | --- | --- | --- | --- |
| 1 | 1 | 36 | 4 | -23 | 50 | 5 | -30 | -33 | 23 | 98 | 12 | -36 | -10 | -7 |
| 2 | 1 | 38 | -18 | -2 | 51 | 7 | -31 | -14 | 19 | 99 | 12 | -29 | -22 | 16 |
| 3 | 1 | 42 | -18 | -2 | 52 | 7 | -36 | -14 | 19 | 100 | 13 | -36 | 4 | 3 |
| 4 | 1 | 30 | 24 | 5 | 53 | 7 | -30 | 7 | 12 | 101 | 13 | -40 | 4 | 5 |
| 5 | 1 | 39 | 2 | -9 | 54 | 7 | -28 | 10 | 18 | 102 | 13 | -36 | 4 | 3 |
| 6 | 1 | 38 | 6 | -4 | 55 | 8 | -33 | 10 | 1 | 103 | 13 | -40 | 4 | 5 |
| 7 | 1 | 37 | 8 | 0 | 56 | 8 | -31 | 12 | 6 | 104 | 13 | -33 | 6 | -11 |
| 8 | 1 | 36 | 10 | 5 | 57 | 8 | -31 | 14 | 10 | 105 | 13 | -33 | 11 | -6 |
| 9 | 1 | 35 | 12 | 11 | 58 | 8 | -30 | 15 | 15 | 106 | 13 | -31 | 17 | 0 |
| 10 | 1 | 38 | -8 | 8 | 59 | 8 | -37 | -6 | -8 | 107 | 13 | -30 | 23 | 6 |
| 11 | 1 | 37 | -10 | 13 | 60 | 8 | -36 | -10 | -2 | 108 | 13 | -32 | -23 | 19 |
| 12 | 1 | 36 | -13 | 19 | 61 | 8 | -35 | -13 | 2 | 109 | 13 | -37 | -23 | 20 |
| 13 | 2 | -30 | 7 | 18 | 62 | 8 | -35 | -15 | 6 | 110 | 13 | -37 | -2 | -8 |
| 14 | 2 | -39 | -1 | -19 | 63 | 8 | -34 | -19 | 11 | 111 | 13 | -35 | -6 | -1 |
| 15 | 2 | -32 | -28 | 6 | 64 | 8 | -34 | -22 | 15 | 112 | 13 | -34 | -11 | 6 |
| 16 | 2 | -38 | -8 | -12 | 65 | 8 | -32 | -25 | 20 | 113 | 13 | -33 | -16 | 13 |
| 17 | 2 | -44 | -8 | -11 | 66 | 8 | -32 | -27 | 24 | 114 | 13 | -31 | -22 | 19 |
| 18 | 2 | -28 | 10 | 13 | 67 | 9 | 36 | 5 | -9 | 115 | 14 | 30 | 14 | 5 |
| 19 | 2 | -27 | 12 | 18 | 68 | 9 | 35 | 9 | -4 | 116 | 14 | 34 | 15 | 7 |
| 20 | 2 | -42 | -6 | -5 | 69 | 9 | 32 | 12 | 1 | 117 | 14 | 38 | 15 | 8 |
| 21 | 2 | -41 | -8 | 0 | 70 | 9 | 31 | 15 | 4 | 118 | 14 | 31 | -21 | 20 |
| 22 | 2 | -40 | -11 | 4 | 71 | 9 | 30 | 19 | 9 | 119 | 14 | 36 | -23 | 20 |
| 23 | 2 | -39 | -14 | 9 | 72 | 9 | 28 | 23 | 14 | 120 | 14 | -37 | -16 | 16 |
| 24 | 2 | -38 | -16 | 14 | 73 | 10 | -34 | -12 | 20 | 121 | 14 | -42 | -16 | 16 |
| 25 | 2 | -37 | -18 | 18 | 74 | 10 | -27 | 14 | 9 | 122 | 15 | -35 | -16 | 14 |
| 26 | 2 | -35 | -21 | 23 | 75 | 10 | -32 | 16 | 8 | 123 | 15 | -40 | -17 | 15 |
| 27 | 3 | 37 | 8 | 2 | 76 | 10 | -37 | 18 | 7 | 124 | 15 | -30 | 12 | 9 |
| 28 | 3 | 36 | 11 | 6 | 77 | 10 | -32 | 10 | -5 | 125 | 15 | -36 | 12 | 12 |
| 29 | 3 | 35 | 14 | 10 | 78 | 10 | -31 | 13 | -1 | 126 | 15 | -37 | 8 | -10 |
| 30 | 3 | 34 | 17 | 14 | 79 | 10 | -31 | 16 | 2 | 127 | 15 | -35 | 11 | -6 |
| 31 | 3 | 31 | 28 | 2 | 80 | 10 | -29 | 20 | 7 | 128 | 15 | -34 | 14 | -3 |
| 32 | 4 | 29 | 17 | 7 | 81 | 10 | -28 | 22 | 11 | 129 | 15 | -33 | 18 | 2 |
| 33 | 4 | 34 | 18 | 8 | 82 | 11 | 40 | -23 | -3 | 130 | 15 | -32 | 21 | 5 |
| 34 | 4 | 38 | 18 | 8 | 83 | 11 | 33 | 7 | -6 | 131 | 15 | -30 | 26 | 9 |
| 35 | 4 | 36 | -22 | 2 | 84 | 11 | 38 | 3 | -7 | 132 | 15 | -39 | -17 | 3 |
| 36 | 4 | 41 | -22 | 2 | 85 | 11 | 37 | 0 | 1 | 133 | 15 | -44 | -17 | 3 |
| 37 | 4 | 40 | 1 | -7 | 86 | 11 | 35 | -4 | 9 | 134 | 16 | -35 | 4 | 5 |
| 38 | 4 | 38 | 5 | -1 | 87 | 11 | 35 | -7 | 17 | 135 | 16 | -39 | 4 | 7 |
| 39 | 4 | 38 | 9 | 3 | 88 | 11 | -35 | 1 | -16 | 136 | 16 | -32 | -21 | 16 |
| 40 | 4 | 37 | 12 | 7 | 89 | 11 | -35 | 6 | -10 | 137 | 16 | -37 | -19 | 18 |
| 41 | 4 | 37 | 16 | 11 | 90 | 11 | -34 | 12 | -4 | 138 | 16 | -36 | -21 | -5 |
| 42 | 5 | -36 | -10 | 15 | 91 | 11 | -33 | 16 | 2 | 139 | 16 | -40 | -22 | -4 |
| 43 | 5 | -41 | -10 | 15 | 92 | 11 | -31 | 22 | 9 | 140 | 16 | -30 | -21 | 13 |
| 44 | 5 | -34 | 1 | -8 | 93 | 12 | -37 | -4 | -8 | 141 | 16 | -29 | -27 | 20 |
| 45 | 5 | -31 | 10 | 6 | 94 | 12 | -36 | -3 | -3 | 142 | 17 | -29 | 22 | 9 |
| 46 | 5 | -30 | 12 | 10 | 95 | 12 | -35 | 0 | 3 | 143 | 17 | -33 | 22 | 9 |
| 47 | 5 | -29 | 16 | 15 | 96 | 12 | -35 | 1 | 9 | 144 | 17 | -35 | 9 | -10 |
| 48 | 5 | -30 | -26 | 15 | 97 | 12 | -34 | 2 | 14 | 145 | 17 | -33 | 13 | -6 |

| No | Subject | X | Y | Z |
| --- | --- | --- | --- | --- |
| 146 | 17 | -31 | 16 | -4 |
| 147 | 17 | -29 | 22 | 1 |
| 148 | 17 | -28 | 25 | 4 |
| 149 | 17 | -31 | 0 | 17 |
| 150 | 18 | 37 | -10 | 15 |
| 151 | 18 | 41 | -10 | 17 |
| 152 | 18 | 43 | -5 | -12 |
| 153 | 18 | 43 | -9 | -6 |
| 154 | 18 | 41 | -13 | 3 |
| 155 | 18 | 39 | -16 | 11 |
| 156 | 18 | 39 | -20 | 18 |
| 157 | 18 | 39 | -23 | -5 |
| 158 | 19 | 37 | -21 | 4 |
| 159 | 19 | 36 | 16 | -12 |
| 160 | 19 | 35 | 18 | -7 |
| 161 | 19 | 34 | 20 | -2 |
| 162 | 19 | 32 | 22 | 2 |
| 163 | 19 | 31 | 25 | 8 |
| 164 | 20 | 33 | -26 | 5 |
| 165 | 20 | 34 | 5 | -12 |
| 166 | 20 | 32 | 10 | -7 |
| 167 | 20 | 31 | 16 | -3 |
| 168 | 20 | 29 | 21 | 1 |
| 169 | 20 | 28 | 27 | 7 |

**Supplementary Table 6.** Anterior cingulate cortex (ACC) and midcingulate cortex (MCC). MNI coordinates for the 81 ACC and MCC contacts.

| No | Subject | X | Y | Z | No | Subject | X | Y | Z |
| --- | --- | --- | --- | --- | --- | --- | --- | --- | --- |
| 1 | 2 | -6 | 35 | 15 | 49 | 13 | -8 | 38 | 9 |
| 2 | 2 | -10 | 35 | 15 | 50 | 13 | -12 | 39 | 10 |
| 3 | 2 | -15 | 35 | 17 | 51 | 13 | -16 | 42 | 13 |
| 4 | 2 | -10 | 13 | 29 | 52 | 13 | -7 | 13 | 38 |
| 5 | 2 | -14 | 14 | 29 | 53 | 13 | -12 | 15 | 39 |
| 6 | 3 | 11 | 33 | -4 | 54 | 14 | 7 | 46 | 14 |
| 7 | 3 | 7 | 26 | 18 | 55 | 14 | 10 | 49 | 17 |
| 8 | 3 | 12 | 26 | 18 | 56 | 14 | 11 | 13 | 26 |
| 9 | 3 | 9 | 20 | 38 | 57 | 14 | 12 | -14 | 35 |
| 10 | 3 | 10 | -5 | 47 | 58 | 14 | 5 | 38 | -6 |
| 11 | 3 | 15 | -5 | 46 | 59 | 14 | 9 | 38 | -4 |
| 12 | 3 | 10 | 30 | -1 | 60 | 14 | 15 | 39 | 0 |
| 13 | 3 | 13 | 33 | 16 | 61 | 14 | -9 | 29 | 25 |
| 14 | 3 | 13 | 34 | 22 | 62 | 14 | -14 | 31 | 26 |
| 15 | 3 | 14 | 35 | 27 | 63 | 14 | -8 | 11 | 40 |
| 16 | 4 | 8 | 45 | 6 | 64 | 14 | -11 | 13 | 42 |
| 17 | 4 | 13 | 45 | 7 | 65 | 15 | -3 | 38 | 6 |
| 18 | 4 | 18 | 46 | 8 | 66 | 15 | -8 | 39 | 10 |
| 19 | 4 | 10 | 33 | 19 | 67 | 15 | -11 | 40 | 12 |
| 20 | 4 | 16 | 33 | 20 | 68 | 15 | -16 | 42 | 15 |
| 21 | 4 | 10 | 10 | 31 | 69 | 15 | -5 | 33 | 29 |
| 22 | 4 | 17 | 42 | 1 | 70 | 15 | -9 | 35 | 30 |
| 23 | 4 | 17 | 43 | 6 | 71 | 15 | -4 | 16 | 37 |
| 24 | 5 | -6 | 30 | 33 | 72 | 15 | -9 | 16 | 40 |
| 25 | 5 | -4 | 33 | 7 | 73 | 15 | -5 | -2 | 41 |
| 26 | 5 | -5 | 8 | 30 | 74 | 15 | -10 | -3 | 42 |
| 27 | 5 | -10 | 8 | 30 | 75 | 17 | -3 | 44 | 26 |
| 28 | 7 | -12 | 43 | 13 | 76 | 17 | -4 | 16 | 41 |
| 29 | 7 | -15 | 43 | 17 | 77 | 17 | -9 | 17 | 44 |
| 30 | 7 | -11 | 36 | -2 | 78 | 17 | -9 | 4 | 32 |
| 31 | 7 | -14 | 37 | 0 | 79 | 17 | -7 | 35 | 9 |
| 32 | 7 | -10 | 23 | 23 | 80 | 17 | -13 | 37 | 11 |
| 33 | 7 | -5 | -4 | 37 | 81 | 17 | -17 | 39 | 12 |
| 34 | 7 | -12 | -4 | 37 |  |  |  |  |  |
| 35 | 7 | -6 | 4 | 44 |  |  |  |  |  |
| 36 | 7 | -11 | 5 | 45 |  |  |  |  |  |
| 37 | 8 | -8 | 35 | 0 |  |  |  |  |  |
| 38 | 9 | 8 | 33 | -5 |  |  |  |  |  |
| 39 | 9 | 8 | 38 | 31 |  |  |  |  |  |
| 40 | 9 | 14 | 23 | 23 |  |  |  |  |  |
| 41 | 10 | -7 | -2 | 39 |  |  |  |  |  |
| 42 | 10 | -12 | -2 | 39 |  |  |  |  |  |
| 43 | 10 | -5 | 18 | 42 |  |  |  |  |  |
| 44 | 10 | -7 | 35 | -4 |  |  |  |  |  |
| 45 | 10 | -12 | 36 | -2 |  |  |  |  |  |
| 46 | 10 | -8 | 27 | 15 |  |  |  |  |  |
| 47 | 10 | -12 | 28 | 17 |  |  |  |  |  |
| 48 | 13 | -3 | 36 | 6 |  |  |  |  |  |

**Supplementary Table 7.** Posterior cingulate cortex (PCC). MNI coordinates for 31 PCC contacts.

| No | Subject | X | Y | Z |
| --- | --- | --- | --- | --- |
| 1 | 1 | 14 | -44 | 29 |
| 2 | 3 | 11 | -46 | 41 |
| 3 | 3 | 16 | -45 | 42 |
| 4 | 9 | 9 | -33 | 53 |
| 5 | 9 | 9 | -46 | 28 |
| 6 | 9 | 14 | -46 | 27 |
| 7 | 11 | 23 | -37 | -7 |
| 8 | 12 | -15 | -52 | -6 |
| 9 | 12 | -1 | -39 | 35 |
| 10 | 12 | -5 | -38 | 36 |
| 11 | 12 | -10 | -38 | 37 |
| 12 | 12 | -16 | -38 | 39 |
| 13 | 12 | -19 | -38 | 40 |
| 14 | 14 | 8 | -15 | 34 |
| 15 | 14 | 17 | -15 | 36 |
| 16 | 14 | 8 | -38 | 27 |
| 17 | 16 | -6 | -35 | 33 |
| 18 | 16 | -11 | -35 | 34 |
| 19 | 16 | -16 | -36 | 34 |
| 20 | 16 | -12 | -61 | 5 |
| 21 | 16 | -17 | -64 | 6 |
| 22 | 16 | -22 | -65 | 7 |
| 23 | 16 | -7 | -49 | 39 |
| 24 | 16 | -10 | -49 | 43 |
| 25 | 18 | 10 | -48 | 13 |
| 26 | 18 | 14 | -49 | 14 |
| 27 | 18 | 10 | -48 | 13 |
| 28 | 18 | 14 | -49 | 14 |
| 29 | 18 | 6 | -42 | 42 |
| 30 | 18 | 11 | -44 | 43 |
| 31 | 18 | 16 | -45 | 43 |

**Supplementary Table 8.** Intraparietal sulcus (IPS). MNI coordinates for the 62 IPS contacts.

| No | Subject | X | Y | Z | No | Subject | X | Y | Z |
| --- | --- | --- | --- | --- | --- | --- | --- | --- | --- |
| 1 | 1 | 38 | -25 | 40 | 49 | 18 | 42 | -54 | 23 |
| 2 | 1 | 42 | -25 | 39 | 50 | 18 | 46 | -54 | 24 |
| 3 | 1 | 31 | -49 | 57 | 51 | 18 | 51 | -55 | 25 |
| 4 | 1 | 35 | -49 | 56 | 52 | 18 | 29 | -49 | 45 |
| 5 | 1 | 40 | -49 | 56 | 53 | 18 | 34 | -50 | 46 |
| 6 | 1 | 44 | -49 | 55 | 54 | 18 | 39 | -51 | 46 |
| 7 | 1 | 33 | -44 | 26 | 55 | 18 | 44 | -52 | 46 |
| 8 | 1 | 37 | -44 | 25 | 56 | 18 | 48 | -53 | 46 |
| 9 | 1 | 33 | -26 | 42 | 57 | 20 | 34 | -32 | 34 |
| 10 | 1 | 33 | -29 | 48 | 58 | 20 | 39 | -32 | 34 |
| 11 | 2 | -38 | -46 | 23 | 59 | 20 | 44 | -32 | 35 |
| 12 | 2 | -33 | -30 | 37 | 60 | 20 | 49 | -33 | 36 |
| 13 | 3 | 30 | -45 | 45 | 61 | 20 | 53 | -33 | 37 |
| 14 | 3 | 35 | -44 | 46 | 62 | 20 | 58 | -32 | 38 |
| 15 | 3 | 39 | -44 | 47 |  |  |  |  |  |
| 16 | 3 | 44 | -45 | 47 |  |  |  |  |  |
| 17 | 5 | -29 | -40 | 33 |  |  |  |  |  |
| 18 | 5 | -29 | -44 | 38 |  |  |  |  |  |
| 19 | 9 | 30 | -34 | 50 |  |  |  |  |  |
| 20 | 9 | 41 | -36 | 49 |  |  |  |  |  |
| 21 | 9 | 46 | -35 | 50 |  |  |  |  |  |
| 22 | 12 | -37 | -36 | 46 |  |  |  |  |  |
| 23 | 12 | -41 | -36 | 47 |  |  |  |  |  |
| 24 | 12 | -24 | -38 | 42 |  |  |  |  |  |
| 25 | 13 | -29 | -38 | 39 |  |  |  |  |  |
| 26 | 13 | -28 | -42 | 45 |  |  |  |  |  |
| 27 | 13 | -27 | -44 | 52 |  |  |  |  |  |
| 28 | 14 | 32 | -37 | 34 |  |  |  |  |  |
| 29 | 14 | 36 | -38 | 35 |  |  |  |  |  |
| 30 | 14 | 40 | -37 | 36 |  |  |  |  |  |
| 31 | 14 | 45 | -37 | 36 |  |  |  |  |  |
| 32 | 14 | 50 | -37 | 37 |  |  |  |  |  |
| 33 | 16 | -30 | -37 | 33 |  |  |  |  |  |
| 34 | 16 | -36 | -37 | 32 |  |  |  |  |  |
| 35 | 16 | -41 | -38 | 32 |  |  |  |  |  |
| 36 | 16 | -46 | -38 | 32 |  |  |  |  |  |
| 37 | 16 | -51 | -38 | 33 |  |  |  |  |  |
| 38 | 16 | -14 | -50 | 47 |  |  |  |  |  |
| 39 | 16 | -18 | -50 | 51 |  |  |  |  |  |
| 40 | 16 | -24 | -44 | 42 |  |  |  |  |  |
| 41 | 16 | -24 | -50 | 49 |  |  |  |  |  |
| 42 | 16 | -24 | -55 | 57 |  |  |  |  |  |
| 43 | 18 | 42 | -54 | 23 |  |  |  |  |  |
| 44 | 18 | 46 | -54 | 24 |  |  |  |  |  |
| 45 | 18 | 51 | -55 | 25 |  |  |  |  |  |
| 46 | 18 | 35 | -33 | 39 |  |  |  |  |  |
| 47 | 18 | 34 | -38 | 46 |  |  |  |  |  |
| 48 | 18 | 32 | -42 | 53 |  |  |  |  |  |

